# *Arabidopsis* spliceosome factor SmD3 modulates immunity to *Pseudomonas syringae* infection

**DOI:** 10.1101/2021.08.09.455611

**Authors:** Anna Golisz, Michal Krzyszton, Monika Stepien, Jakub Dolata, Justyna Piotrowska, Zofia Szweykowska-Kulinska, Artur Jarmolowski, Joanna Kufel

**Affiliations:** Institute of Genetics and Biotechnology, Faculty of Biology, University of Warsaw, Pawinskiego 5a, 02-106 Warsaw, Poland; Department of Gene Expression, Institute of Molecular Biology and Biotechnology, Faculty of Biology, Adam Mickiewicz University, Umultowska 89, 61-614 Poznan, Poland

**Keywords:** alternative splicing, miRNA, PAMPs, plant immunity, *Pst* DC3000, RNA-seq

## Abstract

SmD3 is a core component of the small nuclear ribonucleoprotein (snRNP) that is essential for pre-mRNA splicing. The role of *Arabidopsis* SmD3 in plant immunity was assessed by testing sensitivity of *smd3a* and *smd3b* mutants to *Pseudomonas syringae* pv. *tomato* (*Pst*) DC3000 infection and its pathogenesis effectors flagellin (flg22), EF-Tu (elf18) and coronatine (COR). Both *smd3* mutants exhibited enhanced susceptibility to *Pst* accompanied by marked changes in the expression of key pathogenesis markers. mRNA levels of these factors were also altered upon treatment with *Pseudomonas* effectors. We showed that SmD3-b dysfunction impairs mainly stomatal immunity as a result of defects in stomatal development. Our genome-wide transcriptome analysis of the *smd3b-1* mutant infected with *Pst* revealed that lack of SmD3-b deregulates defense against *Pst* infection at the transcriptional and posttranscriptional levels including defects in splicing and an altered pattern of alternative splicing. Other changes in the *smd3b-1* mutant involved enhanced elf18- and flg22-induced callose deposition, reduction of flg22-triggered production of early ROS and boost of secondary ROS caused by *Pst* infection. Together, our data indicate that SmD3 contributes to the plant immune response possibly via regulation of mRNA splicing of key pathogenesis factors.

## INTRODUCTION

Plants are challenged by numerous phytopathogens such as bacteria, fungi, and viruses (Muthamilarasan and Prasad, 2013). The best-described hemibiotrophic pathogen is *Pseudomonas syringae* pv. *tomato* DC3000 (*Pst* DC3000), which is a widely used gram-negative bacterial model to assess plant-pathogen interactions and the principles governing plant resistance (Xin and He, 2013). Highly virulent *Pst* DC3000 usually enters host tissue through wounds or stomatal apparatus in leaves and multiplies rapidly in susceptible plants, like *Arabidopsis thaliana*. Plants prevent the entry of *P. syringae* by stomatal closure, activation of salicylic acid (SA)-dependent basal defense and callose deposition in the cell wall that creates a physical barrier at pathogen infection sites (Luna *et al*., 2011; Melotto *et al*., 2008). In turn, *Pst* DC3000 produces phytotoxin coronatine (COR) that activates the jasmonic acid (JA) pathway, induces stomatal reopening and inhibits callose deposition to promote virulence (Bari and Jones, 2009; Geng *et al*., 2014; Luna *et al*., 2011; Melotto *et al*., 2008; Zheng *et al*., 2012).

In addition to mechanistic barriers, plants have developed a two-step specialized innate immune system. The first step is mediated by extracellular pathogen- or microbe-associated molecular patterns (PAMPs/MAMPs) that activate pattern recognition receptors (PRRs) on the cell surface, resulting in immune responses called pattern-triggered immunity (PTI) (Dodds and Rathjen, 2010; Li *et al*., 2016; Macho and Zipfel, 2014; Tang *et al*., 2017; Xin and He, 2013). This response includes PAMP-induced stomatal closure, which is mediated by salicylic acid and abscisic acid (ABA) signaling (Cao *et al*., 2011; Lim *et al*., 2015). The second system, effector-triggered immunity (ETI), is induced by resistance proteins (*R*-proteins), which act as intracellular receptors that recognize avirulence (*Avr*) effectors, often leading to localized programmed cell death (Gimenez-Ibanez and Rathjen, 2010; Li *et al*., 2016). The best characterized PRR receptor in *Arabidopsis* is a leucine-rich repeat receptor-like kinase (LRR-RLK) FLS2 (FLAGELLIN SENSITIVE 2), which recognizes the flg22 peptide from bacterial flagellin protein. Flg22 mimics pathogen appearance and causes oxidative stress, callose deposition and ethylene production, leading to the induction of resistance genes (e.g. *PR1* and *PR5* (*PATHOGENESIS-RELATED GENES*), *PAL1* (*PHE AMMONIA LYASE 1*) and *GSTF6* (*GLUTATHIONE S-TRANSFERASE 6*)), but in contrast to the pathogen does not produce the hypersensitive response (HR) type of necrosis (Asai *et al*., 2002; Gomez-Gomez and Boller, 2002; Maleck *et al*., 2000). Another well-known PRR is the receptor kinase EFR (ELONGATION FACTOR Tu RECEPTOR), which recognizes the elf18 peptide of bacterial translation elongation factor EF-Tu. These PRRs initiate immune signalling by heterodimerization with the LRR-RLK family co-receptor BAK1 (BRI1-ASSOCIATED RECEPTOR KINASE) and recruitment of BIK1 (BOTRYTIS-INDUCED KINASE 1) kinase (Chinchilla *et al*., 2007; Couto and Zipfel, 2016; Macho and Zipfel, 2014; Yeh *et al*., 2016).

During bacterial infection changes in gene expression result mostly from transcriptional reprogramming, but adaptation to biotic stress occurs also on a post-transcriptional level, including pre-mRNA splicing, mRNA export and degradation (Birkenbihl *et al*., 2017; Li *et al*., 2016; Motion *et al*., 2015; Staiger *et al*., 2013; Tsuda and Somssich, 2015). A crucial role of splicing in this process is underlined by pathogen response-related phenotypes of mutants defective in alternative splicing (AS) or the contribution of major AS factors, serine/arginine-rich (SR) proteins, and of alternative splicing of some *R*-genes (e.g. *SNC1* (*SUPPRESSOR OF NPR1-1, CONSTITUTIVE 1*) and *RPS4* (*RESISTANCE TO PSEUDOMONAS SYRINGAE 4*)) to pathogen resistance (Zhang and Gassmann, 2007; Zhang *et al*., 2017). Pathogen response can also be modulated by microRNAs (miRNA) that regulate mRNA stability (Li *et al*., 2011; Ruiz-Ferrer and Voinnet, 2009; Weiberg *et al*., 2014; Zhang *et al*., 2011). For example, miR160, miR167, and miR393 are activated by *Pst* infection or flg22 treatment (Navarro *et al*., 2006), whereas plants lacking or overexpressing miR163 show increased resistance or sensitivity to pathogen, respectively (Chow and Ng, 2017).

SmD3 is one of the core proteins of spliceosome small nuclear ribonucleoprotein (snRNP) complex. Sm proteins (B/B′, D1, D2, D3, E, F and G) directly bind small nuclear RNAs (snRNAs) and are crucial for splicing by contacting pre-mRNA as a part of U snRNP (Zhang *et al*., 2001). The *Arabidopsis* SmD3 homologs, SmD3-a and SmD3-b, contain all conserved regions common to SmD3 proteins, including Sm motifs, an RNA binding domain, and C-terminal GRG and RG domains, suggesting that the function of SmD3 in splicing is preserved in *Arabidopsis* (Swaraz *et al*., 2011). Despite the expression of SmD3-a and SmD3-b in all plant tissues, only the *smd3b* null mutants display pleotropic morphological and developmental phenotypes, including delayed flowering time, reduced root growth and defects in leaf and flower morphology. Consistently, *smd3b* mutation exerts a global effect on pre-mRNA splicing and spliceosome assembly. In contrast, *smd3a* knock-outs have no phenotypic alterations, but the double *smd3a/b* mutant is lethal, suggesting that, although SMD3-b is more important, both proteins have redundant functions.

In this study, we investigated the role of *Arabidopsis* SmD3 in response to biotic stress induced by *Pst*. We show that *smd3a* and *smd3b* mutants are oversensitive to pathogen. Consistently, RNA-seq data for *smd3b-1* plants revealed that lack of SmD3-b dysregulates the course of defense against *Pst* infection at the level of transcription and splicing of factors involved in different aspects of immune response. Since disease susceptibility of *smd3b-1* plants was observed only after surface inoculation, it appears that mainly the pre-invasive stage of defense is attenuated in the mutant, probably resulting from changes in expression of stomatal development and movement genes.

## RESULTS

### Lack of *SmD3* affects resistance to *Pst* DC3000

To investigate the function of SmD3-b protein in plant innate immunity we tested the resistance of the *Arabidopsis smd3b* T-DNA insertion mutants, *smd3b-1* (Figure 1a and 1b) and *smd3b-2* (Figure S1a and S1b) to *Pst* infection by spraying. Upon infection, chlorotic and necrotic symptoms were visible at 72 hpi (hours post infection) both in the wild-type (Col-0) and the *smd3b-1* mutant, but were more severe in the mutant (Figure S1c). Bacteria growth after 24 and 72 hpi for both *smd3b-1* and *smd3b-2* plants showed increased pathogen multiplication compared to the wild-type (Figures 1b and S1b). As the effect was weaker for the *smd3b-2* line, we used the *smd3b-1* mutant for further analyses. Although *smd3a* knock-out has no phenotypic consequences (Swaraz *et al*., 2011), we also checked the effect of *Pst* on *smd3a-1* (Figures 1a and 1c) and *smd3a-2* (Figures S1a and S1b) mutants. Both *smd3a* lines exhibited no statistically significant differences compared to the wild-type (Figures 1c and S1b). To assess cellular defense to the pathogen in *smd3b* and *smd3a* plants we investigated changes in mRNA levels of key pathogenesis markers: *PR1*, *PR2*, *PR4*, *PR5, PDF1.2* (*PLANT DEFENSIN 1.2*) and *GSTF6* that are involved in the SA response, and two *JASMONATE-ZIM-DOMAIN PROTEINS JAZ1* and *JAZ9* from the JA pathway, which are induced by coronatine (Barah *et al*., 2013; Demianski *et al*., 2011; Lieberherr *et al*., 2003). Northern blot and RT-qPCR confirmed that these markers were activated after infection in both Col-0 and the mutants, however, this effect was stronger in *smd3b* and *smd3a* plants after 48 and 72 hpi compared to the wild-type (Figures 1d-f, S1d, S1E, S2). Also, accumulation of three major WRKY transcription factors mRNAs (*WRKY46*, *WRKY53*, and *WRKY70*), that are involved in defense response via the SA pathway and modulate systemic acquired resistance (SAR) (Wang *et al*., 2006), was more prominent in the *smd3b-1* mutant following infection (Figures 1f and S1e). In contrast, expression of other pathogen response-related factors, *SGT1* (*SALICYLIC ACID GLUCOSYLTRANSFERASE 1*)*, NPR1* (*NONEXPRESSER OF PR GENES 1*)*, NPR3* (*NPR1-LIKE PROTEIN 3*) was not significantly altered in the mutant (Figure S1e). Together, these results provide evidence that lack of SmD3 protein dysregulates the response to *Pst* DC3000 infection making plants more susceptible to bacteria.

**Figure 1.**
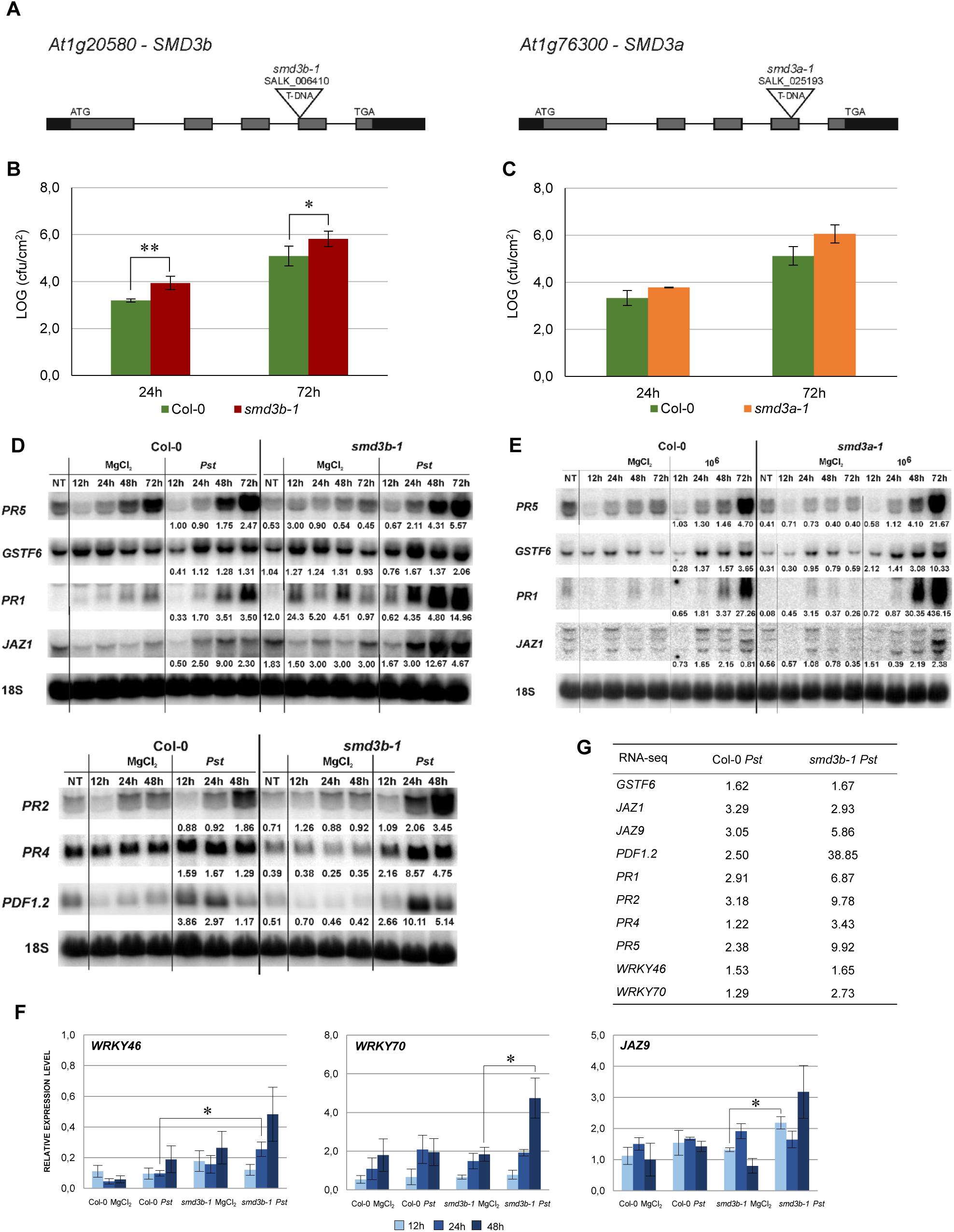
The *smd3b-1* and *smd3a-1* mutants are susceptible to *Pseudomonas syringae* pv. *tomato* DC3000 infection. (a) Structure of the *AtSMD3-B* (*At1g20580*) and *AtSMD3-A* (*At1g76300*) genes. Exons are represented by grey bars, UTRs by black bars and localization of T-DNA insertions are indicated. (b, c) Growth of *Pst* DC3000 after 24 and 72 hpi in Col-0 and the *smd3b-1* (b) or *smd3a-1* (c) mutant. For each time point leaf discs were collected from 5 plants. Results are mean of four independent experiments and error bars represent SEM; *P < 0.05; ** P < 0.01 (Student’s t-test). (d, e) Northern blot analysis of factors involved in response to *Pst* DC3000. Samples were collected from non-treated (NT), control (MgCl_2_) and infected (*Pst*) Col-0 and *smd3b-1* (d) or *smd3a-1* (e) plants at indicated time points. Numbers represent the ratio of transcript level in *Pst-*treated Col-0 and mutants relative to control and normalized to 18S rRNA loading control. Experiments were repeated at least three times; representative blots are shown. (f) RT-qPCR analysis of selected pathogen response genes in *smd3b-1*. Mean values ±SEM were obtained from three independent experiments; *P < 0.05; ** P < 0.01 (Student’s t-test). *UBC9* mRNA was used as a reference. (g) The expression levels of selected pathogen response genes in Col-0 and the *smd3b-1* mutant based on RNA-seq analysis. Numbers represent fold change.

Since SmD3-b is a core component of the snRNP complex, we tested whether a mutant in another Sm protein, SmD1-b, has a similar impact on plant immunity. Bacteria growth assay showed that *smd1b* plants were significantly more sensitive to *Pst* compared to the wild-type, with a similar level of pathogen proliferation as observed for *smd3b* and *smd3a* (Figures 1b, 1c, S1b, S3a). Moreover, as was the case for *smd3b* and *smd3a* lines, activation of key pathogenesis markers, *PR1*, *PR5*, *GSTF6* and *JAZ1*, was also stronger in *smd1b* plants than in Col-0 (Figure S3b). These results reinforce the notion that Sm proteins, most likely as a spliceosomal complex, contribute to shaping the scope of pathogen response signaling.

### The impact of SmD3-b on gene expression in response to *Pst* infection

To estimate the impact of *smd3b-1* mutation on the cell transcriptome under normal conditions and during infection, we sequenced RNA samples from the 6-week-old mutant and Col-0 control and plants treated with *Pst*. Analysis of differential gene expression revealed a significant number of affected genes (DESeq2, FDR (false discovery rate) <0.05; Figure S4a, Dataset S1) between wild-type and the mutant and among treatments. These results show that both *Pst* infection and lack of SmD3-b have profound effects on *Arabidopsis* gene expression. RNA-seq data also confirmed changes in mRNA levels as measured by northern and RT-qPCR, except for *JAZ1* that was downregulated in RNA-seq (see Figure 1g). Principal component analysis (PCA) attested that sequencing data created four coherent groups of biological replicas (Figure S4b). Sets of affected genes showed strong overlaps when compared using an odds ratio statistical test (Figure 2a; all odds ratio > 2.5; GeneOverlap R package (Shen and Sinai, 2013)). Similarity of the lists of genes with changed expression was not limited to sets representing response to infection or impact of *smd3b-1* mutation, but there was also some overlap between genes with expression altered by bacteria and lack of SmD3-b. This supports the notion that *smd3b-1* mutation impacts expression of a subset of genes whose mRNA levels normally change during bacterial attack. Analysis of enriched gene ontology (GO) terms showed that affected mRNAs are related to specific categories that are often common between sets (Figure 2b and Dataset S2). As expected, bacterial treatment upregulated expression of genes involved in defense response and immune system processes in both Col-0 and the mutant. However, enrichment of genes in defense-related GO terms was clearly less prominent in *smd3b-1* than in wild-type plants (Figure 2b).

**Figure 2.**
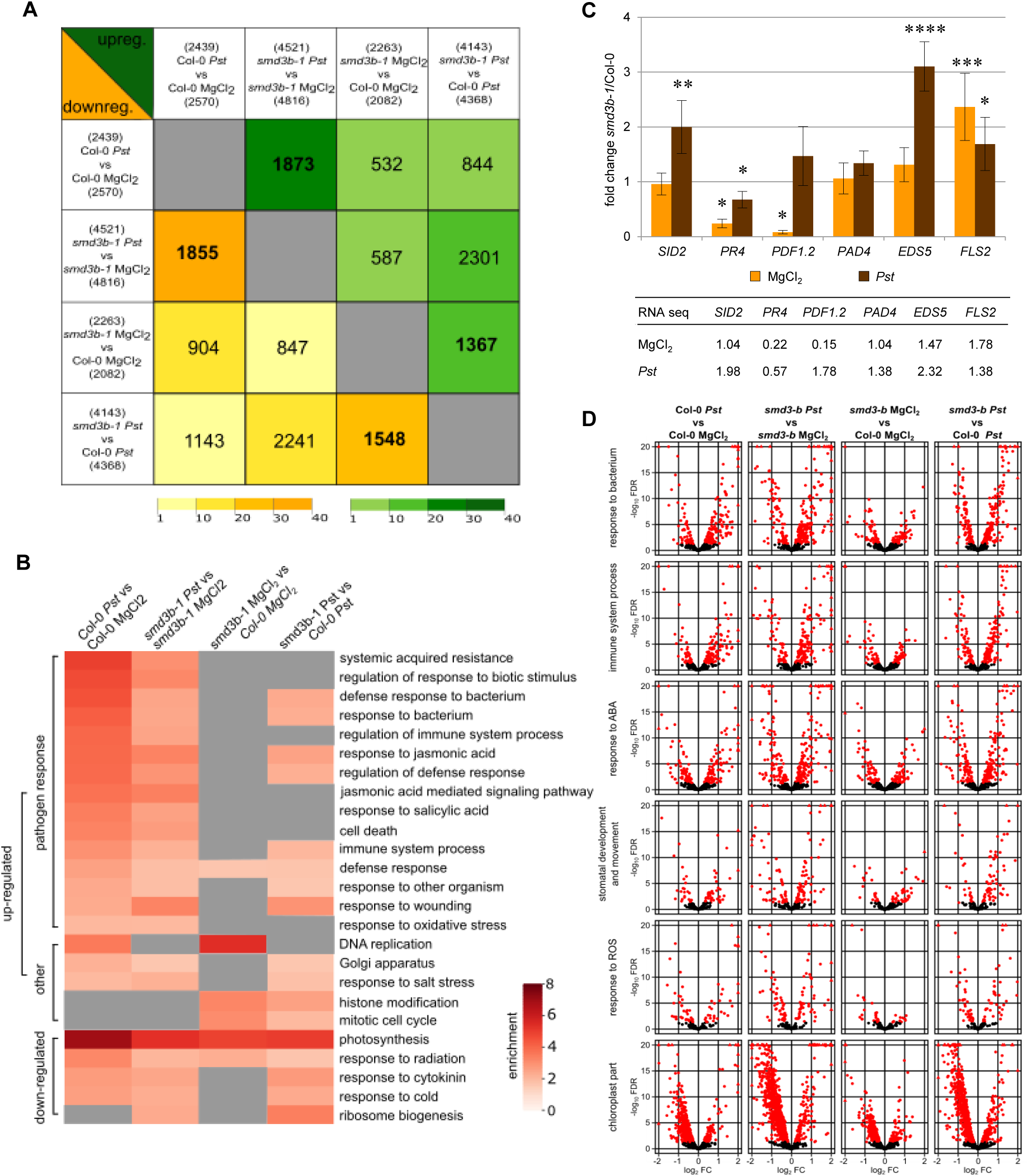
Genes affected by *Pst*-infection and lack of SmD3-b. (a) Affected genes show high overlap between different comparisons. Comparison of genes with changed expression in RNA-seq data for control (MgCl_2_) and infected (48 hpi; *Pst*) Col-0 and *smd3b-1* plants using odds ratio statistical test, which represents the strength of association between the two lists (odds ratio ≤ 1, no association; odds ratio >> 1, strong association). Number of genes with changed expression is shown above (upregulated) and below (downregulated) headings of each comparison; number of overlapping genes is shown in each cell for upregulated (green triangle) and downregulated (yellow triangle) genes, respectively. Colour gradients mark for odds ratio calculated with GeneOverlap R package. (b) Genes with changed expression (FDR < 0.05) show enrichment in several GO categories (shown are only chosen categories, all results are in Dataset S2). (c) Comparison of expression changes for selected mRNAs by RT-qPCR and RNA-seq. Plot show fold changes of transcript levels in *smd3b-1* vs. Col-0 in control condition (MgCl_2_) and 48 hpi (*Pst*). RT-qPCR values represent mean of three independent biological replicates with error bars showing standard deviations (*SD*); \**P* < 0.05; \*\**P* < 0.01; \*\*\**P* < 0.001 (t-test). *UBC9* mRNA was used as a reference. Table presents analogous results from RNA-seq. (d) Bacterial infection and the *smd3b-1* mutation impact gene expression in selected categories. Volcano plots (-log_10_FDR as a function of log_2_FC) for genes implicated in response to bacterium, immune system process, response to abscisic acid (ABA), stomatal development and movement, response to ROS and chloroplast part. Red dots and triangles mark genes with significant changes.

The *smd3b-1* mutant showed altered expression of several genes from the defense response category, either in control or post-infection conditions (Dataset S3, Figures 2d and S5). Notably, a number of genes encoding key pathogenesis-related genes *PR3*, *PR4* and *PR5*, and plant defensin genes, *PDF1.2* and *PDF1.3*, as well as JA- and COR-induced *NATA1* (*N-ACETYLTRANSFERASE ACTIVITY 1*) were markedly downregulated in the mutant in control conditions, but became strongly activated at later time points following *Pst* treatment (Dataset S3, see Figures 1d and 1f). Similar effect was observed for genes of pathogenesis regulatory transcription factors *ANAC019, ANAC055* and *ANAC072* (*NAC DOMAIN CONTAINING PROTEIN*), and *WRKY70*. Another interesting observation in plants lacking SmD3-b was altered expression of factors that regulate BIK1 turnover, namely upregulation of *CPK28* (*CALCIUM-DEPENDENT PROTEIN KINASE 28*), *IRR* (*IMMUNOREGULATORY RNA-BINDING PROTEIN*) and *PERK1/2* (*PEP RECEPTORS*) genes, and downregulation of *PUB26* (*PLANT U-BOX 25/26*) E3 ligase (Dataset S3).

Antibacterial defense is regulated by biotic stress hormones SA, JA and ABA that are involved in controlling stomatal movement, MAPK signaling, generation of reactive oxygen species and stimulation of callose deposition. The expression of several factors of the SA-JA and ABA signaling pathways was altered in the *smd3b-1* mutant in control conditions or during *Pst* infection (Dataset S3, Figures 2b and S6). Among the most important changes was elevated expression of protein phosphatase *PP2C/HAI1* (*HIGHLY ABA-INDUCED PP2C 1*) in both control and post-infection conditions. PP2C/HAI1, induced by ABA and COR upon *Pst* infection, dephosphorylates MPK3 and MPK6 kinases leading to their inactivation and immune suppression (Mine *et al*., 2018). In turn, genes involved in SA synthesis and signaling were more strongly induced by *Pst* infection in *smd3b-1*. This concerns for example SA-synthesis genes *EDS5* (*ENHANCED DISEASE SUSCEPTIBILITY 5*)*, PBS3* (*AVRPPHB SUSCEPTIBLE 3*) and *SID2/ICS1* (*SALICYLIC ACID INDUCTION DEFICIENT 2/ISOCHORISMATE SYNTHASE 1*), N-hydroxy pipecolic acid (NHP)-synthesis genes *FMO1* (*FLAVIN-DEPENDENT MONOOXYGENASE 1*) and *ALD1* (*AGD2-LIKE DEFENSE RESPONSE PROTEIN 1*) as well as SA methyltransferase and methyl esterase genes *BSMT1* (*BA/SA CARBOXYL METHYLTRANSFERASE 1*) and *MES9* (*METHYL ESTERASE 9*). Other relevant differences between *smd3b-1* and Col-0 plants related to the hormonal crosstalk include altered activation of ABA biosynthesis gene *NCED3* (*NINE-CIS-EPOXYCAROTENOID DIOXYGENASE 3*), upregulated expression of JA biosynthesis enzyme *LOX2* (*LIPOXYGENASE 2*) and negative transcriptional repressors of the JA-responsive genes, *JAZ1*, *JAZ5* and *JAZ9*.

Changes in gene expression were confirmed by RT-qPCR for six selected defense response-related genes (Figure 2c). As seen previously (Figures 1d and 1f), *smd3b-1* mutation resulted in a significant decrease of *PDF1.2* and *PR4* mRNAs under normal conditions, whereas expression of *FLS2*, which is required for the perception of PAMP flagellin, was markedly increased. In turn, after *Pst* treatment *SID2/ICS1*, *PR4* and *EDS5* showed significant upregulation in the mutant. These results, as well as our northern and RT-qPCR analyses (Figures 1d, 1f, S1d, S1e), were consistent with RNA-seq (Figure 1g) so altered expression of pathogenesis markers in the *smd3b-1* mutant before and after *Pst* treatment was confirmed by three different methods. These observations suggest that the *smd3b-1* mutant shows perturbations in defense response to bacteria, including regulation of the immune system and biotic stress hormones. Both *Pst* treatment and *smd3b-1* mutation affected expression of genes involved in other numerous processes including histone modification, DNA replication, cell cycle, Golgi apparatus, chloroplast stroma and thylakoid. Another interesting observation was that pathogen treatment of the mutant decreased expression of genes involved in photosynthesis, chloroplast activity, and ribosome biogenesis and function (Figures 2d and S5). Northern hybridizations using probes located downstream of 18S and 5.8S rRNA revealed moderately altered level of 35S and 25SA/B rRNA precursors in the *smd3b-1* mutant upon *Pst* infection, confirming that pre-rRNA processing is indeed affected (Figure S6). Still, the bases of these effects or their implications for bacterial infection are unclear.

Differences between *smd3b-1* and wild-type plants in expression of genes related to pathogenesis in control and post-infection conditions are well illustrated by clustering analysis (Figure S7). Of special interest is a large number of genes in response to bacterium and immune system GO terms that upon *Pst* infection are strongly upregulated in Col-0 but have a much lower expression in the mutant (clusters 1, 2, Figure S7a). Another class represents genes in the same GO terms with *Pst*-induced expression in Col-0 that become even more highly activated in *smd3b-1* (cluster 3, Figure S7a). In turn, many genes that are downregulated in Col-0 in response to *Pst* often have decreased expression in *smd3b-1* plants already in control conditions and are affected by the pathogen to a lesser extent (clusters 1 in upper panel and 1, 2 in lower panel, Figure S7b). The latter behaviour is also observed for many photosynthesis- and chloroplast-associated genes (clusters 1, 2, Figure S7c). A general *Pst*-mediated suppression of these genes reflects the central role of chloroplasts in plant immunity as a major site for production of reactive oxygen species (ROS) and defense-related hormones SA, JA, and ABA (Lewis *et al*., 2015; Lu and Yao, 2018; Serrano *et al*., 2016; de Torres Zabala *et al*., 2015).

We also analysed global changes in mRNA alternative splicing using rMATS (Shen *et al*., 2014), which allowed identification of AS events significantly altered during *Pst* infection or by the *smd3b-1* mutation (FDR < 0.05; ΔPSI (percent spliced-in) > 0.05; Table 1, Dataset S4). The highest number of affected splicing events was observed after *Pst* treatment in the mutant compared to the wild-type. In turn, comparing these two lines in control conditions identified fewer changes in the AS pattern, whereas *Pst* infection alone generated far less AS events in both wild-type and mutant plants. Based on the number of events and affected genes, we conclude that lack of SmD3-b protein has a greater impact on the splicing pattern than response to pathogenic bacteria. Nonetheless, splicing deficiency resulting from Smd3-b dysfunction is further exacerbated by pathogen infection. A high number of differential events for each comparison (from 25.3 to 33.4% depending on the set) was novel according to AtRDT2 annotation (Zhang *et al*., 2017). It appears that retained introns (RI) represented the most common AS events (from 55.3 to 71.8%), while alternative 5’ and 3’ spliced sites (A5/A3) or skipped exons (SE) were much less numerous (Table 1). Further assessment revealed that from 45.9 to 65.7% of splicing events and from 26.8 to 41.8% of genes with splicing events were unique when compared to other sets (Figure S8a). As expected, lack of the core snRNP protein SmD3-b destabilizes U1, U2 and U4 snRNAs (Dataset S3 and Figure S8b), supporting a general splicing defect in *smd3b* plants (Swaraz *et al*., 2011). Differential splicing events for six genes were visualised using Sashimi plots and verified by RT-qPCR (Figure S9a). All of these genes showed changes in the splicing pattern due to *smd3b-1* mutation, however, splicing of *At5g20250* and *At5g57630* was also altered in wild-type plants during *Pst* infection. Treatment with flg22 had a similar impact on the splicing pattern in the mutant as the pathogen (Figure S9b). These results suggest that both early PAMP-induced and later pathogen-triggered stages of infection affect SMD3-regulated splicing events. Among genes having significantly affected AS events in the mutant we found several genes involved in pathogen response, including *MAP4K*, *MPK17*, *WRKY60*, *CRK4*, *SUA, SNC1*, *RIN2*, *NINJA, BAK1*, *CPK28*, *SNC4*, *At3g44400*, *PRX34*, *AGG1*, *MEKK3*, *CRK6*, *XLG2* and *TGA2* (Figures 3 and S9c, Table S1, Dataset S4).

**Figure 3.**
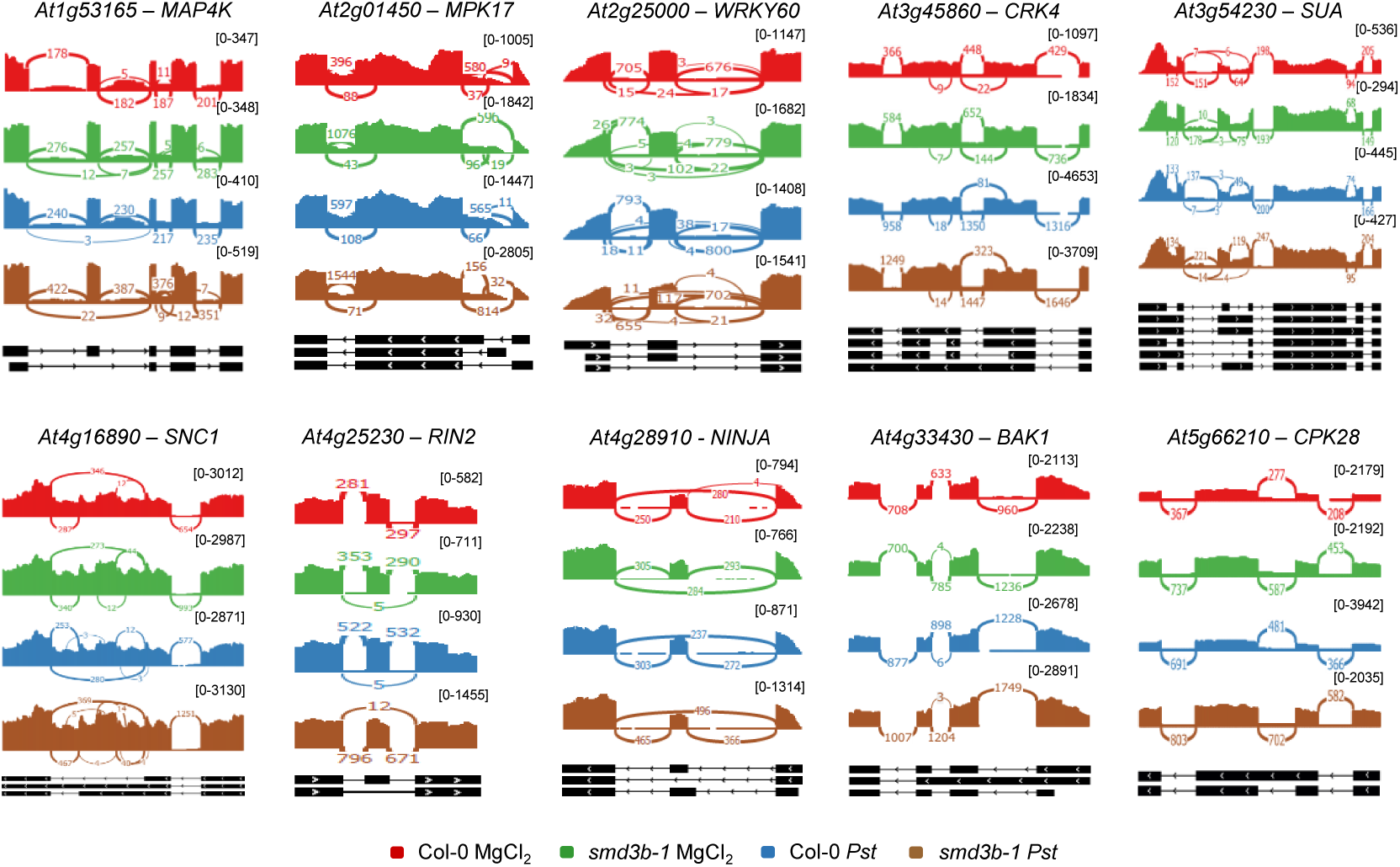
Both *Pst* infection and *smd3b-1* mutation affects alternative splicing (AS) events. Analysis of AS events for selected genes in control (MgCl_2_) and infected (48 hpi; *Pst*) Col-0 and *smd3b-1* plants. AS events are significantly altered for genes involved in pathogen response. Sashimi plots were created from RNA-seq data using Integrative Genomics Viewer (IGV). The numbers in brackets are the range on the bar graph. Note differences in scales for each sashimi plot.

**Table 1.** Alternative splicing events and genes significantly altered by infection or *smd3b-1* mutation.

|  | Col-0 *Pst*<br>vs<br>Col-0 MgCl<sub>2</sub> | *smd3b-1 Pst*<br>vs<br>*smd3b-1* MgCl<sub>2</sub> | *smd3b-1* MgCl<sub>2</sub><br>vs<br>Col-0 MgCl<sub>2</sub> | *smd3b-1 Pst*<br>vs<br>Col-0 *Pst* |
| --- | --- | --- | --- | --- |
| genes | 1010 (217) | 775 (234) | 1685 (384) | 1955 (495) |
| events | 1261 (319) | 961 (320) | 2524 (705) | 3025 (1012) |
| A3 | 158 (68) | 118 (71) | 336 (175) | 344 (195) |
| A5 | 104 (1) | 108 (4) | 389 (22) | 326 (21) |
| RI | 906 (223) | 640 (207) | 1396 (378) | 1844 (603) |
| SE | 82 (17) | 79 (28) | 366 (99) | 470 (159) |
| MX | 11 (10) | 16 (10) | 37 (31) | 41 (34) |

Abbreviation: A3 – alternative 3' splice site; A5 – alternative 5' splice site; RI – retained intron; SE – skipping exon; MX – mutually exclusive exons. The numbers in brackets represent novel alternative splicing events and genes.

Global analysis of the transcriptome showed that *smd3b-1* mutation affected mRNA levels and splicing, including alternative splicing, both in normal conditions and during *Pst* infection. Since these changes concern several pathogenesis factors we conclude that this outcome directly or indirectly impacts the response to pathogen attack.

### Lack of SmD3 dysregulates the response to biotic effectors

PAMPs directly activate the innate immune response in plants. Flg22 and elf18 are recognized by PRR receptors, which induce a pattern-triggered immunity response mediated by receptor-like kinases FRK1 (FLG22-INDUCED RECEPTOR-LIKE KINASE 1) and BAK1. These proteins activate the MAP kinase cascade, leading to the expression of defense genes, such as *WRKY* transcription factors. In turn, COR stimulates JA-signalling and in consequence suppresses SA-dependent defense (Brooks *et al*., 2005).

To investigate the mode of action of PAMP-induced pathways in the *smd3b-1* mutant, we treated 14-day-old seedlings with flg22, elf18, or COR and examined the level of selected mRNAs involved in biotic stress responses by northern blot (Figure 4a). In accordance with their biological activities, flg22 and elf18 induced the expression of *FRK1*, *BAK1,* and *WRKY29*, whereas COR treatment activated *JAZ1*, *JAZ3*, *MYC2* (*JASMONATE INSENSITIVE 1*), and *VSP2* (*VEGETATIVE STORAGE PROTEIN 2*) components of the MYC branch of the JA pathway in both wild-type and mutant plants compared to control conditions. These results confirmed the specificity of flg22/elf18- and COR-mediated pathways, as their respective components were not affected by unrelated PAMPs. The pattern of pathogen markers in the *smd3b-1* mutant differed from that in wild-type plants, especially for *CORI3* (*CORONATINE INDUCED 1*), *PR1*, *PR2* and *FRK1*, which showed elevated basal expression, suggesting constitutive activation of stress response genes. Also, some transcripts were induced by PAMPs to varying extents in the mutant, e.g. activation of *BAK1* was stronger and *FRK1* weaker following flg22 or elf18 treatment, the levels of *GSTF6* and *JAZ1* were upregulated by flg22, while COR-triggered accumulation was lower for *CORI3* and slightly higher for *MYC2* (Figure 4a).

**Figure 4.**
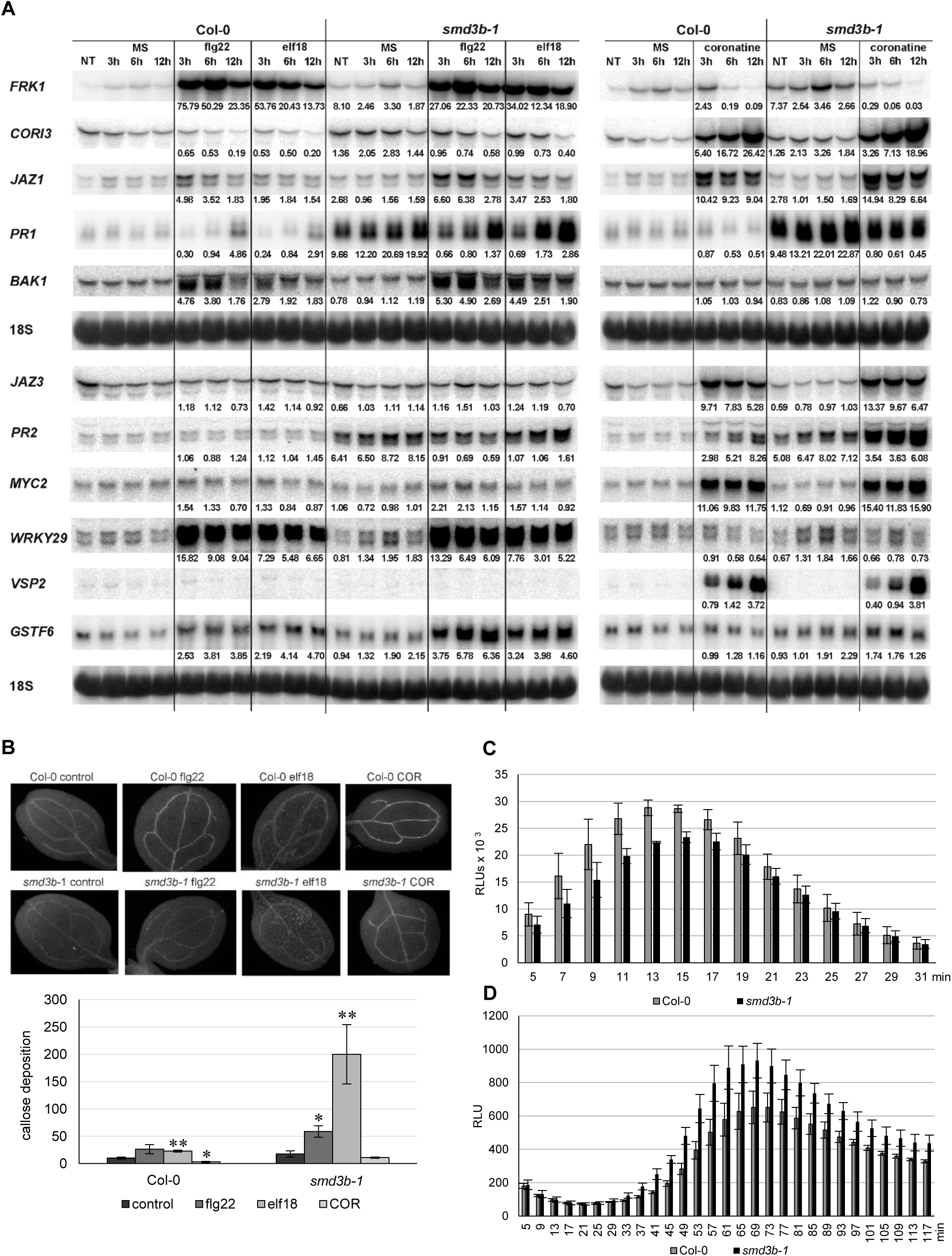
PAMP-induced expression of pathogenesis markers, callose deposition and production of ROS is altered in the *smd3b-1* mutant. (a) Northern blot analysis of factors involved in PAMP response. Samples were collected at indicated time points from non-treated (NT) 14-day-old seedlings, treated with MS (control) or 100 nM of flg22, elf18 and COR. The ratio of transcript level in treated Col-0 and *smd3b-1* relative to control (MS) is shown as main numbers, while the ratio of control *smd3b-1* vs. Col-0 is given in parentheses. Values were normalized to 18S rRNA loading control. Experiments were repeated four times; representative blots are shown. (b) One-week-old plants were treated with MS (control) or 1µM of flg22, elf18 and COR. Callose formation was visualized by aniline blue staining and epifluorescence microscopy and quantified using ImageJ software from digital photographs as a number of local maxima specified by the average of RGB coloured pixels (callose intensity) in plant material. Bars represent mean of three independent biological replicates with error bars showing *SD*; \**P* < 0.05; \*\**P* < 0.01 (Student’s t-test). Representative pictures are shown. (c, d) ROS production in response to 100 nM flg22 (c) or *Pst* (d) treatment in leaf discs from 6-week-old Col-0 and *smd3b-1* plants. Bars represent mean of three independent biological replicates with error bars showing *SD*; \**P* < 0.05; \*\**P* < 0.01 (Student’s t-test). Luminescence is in Relative Light Units (RLUs).

These data confirm that flg22/elf18- and COR-induced responses represent separate pathways and show that SmD3-b moderately affects PAMP-triggered activation of early- or/and late-responsive genes involved in plant innate immunity. They are also consistent with the contribution of COR to pathogenesis by suppressing basal defense-associated genes (Thilmony *et al*., 2006).

Another important indicator of PTI is PAMP-induced deposition of callose in cotyledons or leaves. To test effects of *smd3b-1* mutation on callose accumulation, the mutant and wild-type seedlings were treated with flg22, elf18, or COR effectors and examined by microscopy 24 h after treatment (Figure 4b). Quantification of the callose signal revealed that elf18- and flg22-induced deposition of callose was significantly higher in the *smd3b-1* leaves than in Col-0. As shown previously, callose production was suppressed by COR treatment (Geng *et al*., 2012). This result, suggesting that lack of SmD3-b somehow promotes accumulation of callose, is rather counter-intuitive as callose is supposed to reinforce the cell wall against pathogen entry (Ellinger *et al*., 2013). However, callose and pathogen resistance is modulated by SA signaling and is affected by several factors, such as growth and stress conditions, thus callose deposition does not always match the activity of plant immunity (Luna *et al*., 2011; Nishimura *et al*., 2003).

*Pst* infection through PAMPs also elicits production of reactive oxygen species. The primary, low-amplitude, apoplastic ROS, dependent on cell wall peroxidases and plasma membrane NADPH oxidases, triggers PTI-dependent basal antimicrobial defense (Daudi *et al*., 2012; Jwa and Hwang, 2017; Shapiguzov *et al*., 2012). To test whether SmD3 is required for PTI activation, we measured the levels of flg22-triggered H_2_O_2_ in wild-type and *smd3b-1* plants. A luminol-based assay for leaves treated with flg22 revealed that ROS accumulation was significantly reduced in the mutant compared to Col-0 (Figure 4c). This effect may be due to attenuation of RBOHD (Respiratory Burst Oxidase Homolog D) activity. RBOHD is the major ROS-generating plasma membrane NADPH oxidase and is regulated by phosphorylation and ubiquitination (Kadota *et al*., 2015; Lee *et al*., 2020). One of the the RBOHD-phosphorylating kinases, receptor-like cytoplasmic kinase PBL13 (PBS1-like kinase 13), acts as a negative regulator of RBOHD stability and activity (Lee *et al*., 2020). Expression of *PBL13* is markedly upregulated in the absence of SmdD3-b (S3 Dataset) and this may lead to destabilization of RBOHD and decreased ROS production. Next, we also checked the production of the secondary, high-amplitude, chloroplastic ROS after pathogen treatment and found out that, in contrast to the apoplastic ROS, a markedly higher intracellular ROS was generated in the *smd3b-1* mutant (Figure 4d). These results suggest that while a weaker burst of apoplastic ROS in the absence of SmD3-b may result in a less effective inhibition of pathogen multiplication, a stronger accumulation of intracellular ROS possibly reinforces plant stress response by activation of defense-related genes.

Different stress conditions that involve ROS production evoke endonucleolytic cleavage of tRNA molecules at the anticodon loop (Thompson *et al*., 2008). tRNA fragments (tRFs) may contribute to translation inhibition during microbial attack or act as stress response signaling molecules (Schimmel, 2018). To determine whether biotic stress in *Arabidopsis* also results in tRNA fragmentation, we checked decay intermediates for a few tRNAs in Col-0 and *smd3b-1* plants following treatment with *Pst* at 24, 48, and 72 hpi (Figure S10). As expected, bacterial infection led to accumulation of shorter RNA fragments indicative of tRNA cleavage. Interestingly, the amount of tRFs was increased in the mutant, in line with a higher ROS level. It is therefore possible that this outcome reflects a compromised defense of the mutant towards the pathogen.

Taken together, these results show that SmD3 modulates several aspects of early PTI and late ETI responses.

### Lack of SmD3-b affects pathogen entry

Sensitivity to the pathogen may also arise due to alterations in stomatal functioning since stomata play an important role in plant immunity as a major entryway for bacteria (Cao *et al*., 2011; Lim *et al*., 2015; Melotto *et al*., 2006). RNA-seq data revealed that a large number of genes that are involved in stomatal development, movement or dynamics were significantly changed in the *smd3b-1* mutant, in both control and post-infection conditions (Figure 2d and Dataset S3). These included regulators of stomatal density and patterning (e.g. *STOMAGEN* (*EPFL9, EPIDERMAL PATTERNING FACTOR-LIKE 9*)*, EPF2* (*EPIDERMAL PATTERNING FACTOR 2*)*, TMM* (*TOO MANY MOUTHS*)*, ER* (*ERECTA*) and *ERL1* (*ERECTA LIKE 1*), ABA-induced stomatal closure (e.g. PP2C phosphates *ABI1* (*ABA-INSENSITIVE 1*), *ABI2* and *HAB1* (*HOMOLOGY TO ABI1 1*), ubiquitin E3 ligase *CHYR1*/*RZPF34* (*CHY ZINC-FINGER AND RING PROTEIN 1/RING ZINC-FINGER PROTEIN 34*), *GHR1* (*GUARD CELL HYDROGEN PEROXIDE-RESISTANT 1*), *SLAC1* (*SLOW ANION CHANNEL-ASSOCIATED 1*), *SIF2* (*STRESS INDUCED FACTOR 2*), and stomatal reopening (e.g. NAC transcription factors *ANAC019, ANAC055* and *ANAC072,* SA synthesis and modification enzymes *SID2*/*ICS1* and *BSMT1*). Finally, altered were also the levels of *LecRK-V.5* and *LecRK-VI.2* (*LEGUME-LIKE LECTIN RECEPTOR KINASES*) that act as negative and positive regulators of stomatal immunity, respectively (Dataset S3) (Arnaud and Hwang, 2015; Pillitteri and Torii, 2012; Simmons and Bergmann, 2016; Zhang *et al*., 2020). We therefore checked the state of stomatal density and aperture in wild-type and *smd3b-1* plants and observed that both were significantly increased in the mutant (Figures 5a and 5b). Such features may enable faster entry and facilitate proliferation of bacteria in *smd3b-1* plants, and as a consequence increase sensitivity to pathogen. To confirm this possibility we used a different infection method, i.e. syringe infiltration, in which the pathogen is not delivered into the leaf tissue *via* stomata as in a natural situation but directly to the apoplastic space. The results showed that, as opposed to surface inoculation by spraying, *smd3b-1* plants were no longer sensitive to *Pst* after infiltration inoculation (Figure 5c). Moreover, also in stark contrast to spraying, induction of key pathogenesis markers (*PR1*, *PR2*, *PR4*, *PR5*, *PDF1*.2 and *GSTF6*) after 48hpi was similar or even weaker in the *smd3b-1* mutant compared to Col-0 (Figure 5d). This result is consistent with observations that *Pst*-induced changes in gene expression in the mutant are more prominent than these triggered by PAMPs (see Figures 1d, 1e and 4a). Such an outcome may reflect a situation when pre-invasive stage of defense response, i.e. pathogen entry, is affected in the mutant. Since pathogen entry to the apoplastic space is restricted by PAMP-induced stomatal closure mediated by ABA signaling in guard cells, we next tested stomatal movement following ABA treatment. There was virtually no difference in ABA-stimulated stomatal closure between wild-type and *smd3b-1* plants (Figure S11), indicating that this aspect of defense signaling was not impaired in the mutant. Still, the higher number of stomata and their larger pores in the *smd3b-1* probably lead to initial more unrestrained pathogen entry. Together these data suggest that proper functioning of SmD3b contributes to establishing effective stomatal immunity.

**Figure 5.**
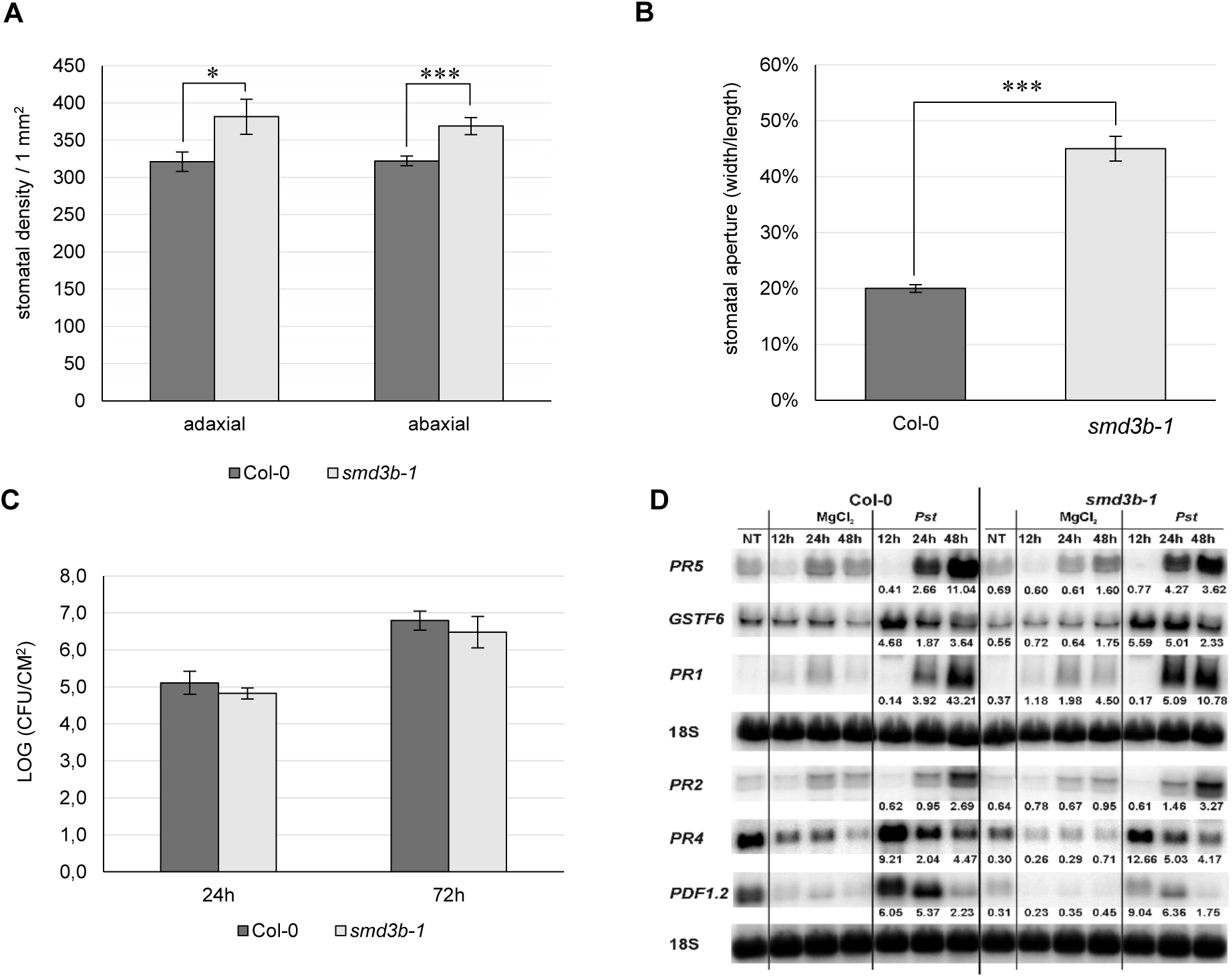
SmD3 contributes to stomatal immunity. (a, b) Stomatal density and aperture of Col-0 and *smd3b-1* plants. Comparison of the stomatal density on the abaxial and adaxial leaf side of Col-0 and *smd3b-1*, shown as number of stomata per 1 mm^2^ of leaf surface (a). Stomatal aperture in Col-0 and *smd3b-1* (b). The aperture on the abaxial leaf side was calculated as stomata width to length to ratio and expressed as percentage. (c) The *smd3b-1* mutant is not sensitive to *Pst* after infiltration inoculation. Growth of *Pst* DC3000 at 24 and 72 hpi after syringe infiltration in Col-0 and the *smd3b-1* mutant. For each time point leaf discs were collected from 8 plants. Results are mean of two independent experiments and error bars represent SD. (d) Northern blot analysis of factors involved in pathogen response. Samples were collected from non-treated (NT), control (MgCl_2_) and injected (*Pst*) Col-0 and *smd3b-1* plants at indicated time points. Numbers represent transcript level in *Pst*-treated Col-0 and the *smd3b-1* relative to control and normalized to 18S rRNA loading control.

### SmD3-b modifies levels of pri-miRNA and mature miRNA upon *Pst* infection

Since plant miRNAs are differentially expressed during pathogen infection and may contribute to the regulation of plant immunity we evaluated changes in pri-miRNA and miRNA levels in *Pst*-infected 6-week-old *smd3b-1* and Col-0 plants (48 hpi). RT-qPCR analysis showed significant changes for 9 out of 14 tested pri-miRNAs (Figures 6a and S12). In the control condition, the level of pri-miR156A and pri-miR403 was reduced in the mutant compared to the wild-type, whereas expression of pri-miR161, pri-miR171A, pri-miR171C, and pri-miR393B was downregulated following pathogen treatment. In contrast, accumulation of pri-miR163 and pri-miR393A was increased after *Pst* infection in both mutant and wild-type plants. The expression of corresponding mature miRNAs was analysed by northern blot (Figure 6b). We observed that after *Pst* treatment miR163, miR393, and miR319 were upregulated to a lesser extent in the *smd3b-1* than in wild-type plants. We also tested the level of *FAMT* mRNA, which is the one of the targets of miR163 and has a direct role in pathogen response (Chow and Ng, 2017). In agreement with accumulation of miR163, the induction of *FAMT* mRNA 48 hpi was significantly stronger in the mutant compared to the wild-type (Figure 6c). These results suggest a correlation between expression of miRNAs and their targets to modulate defense response against pathogen.

**Figure 6.**
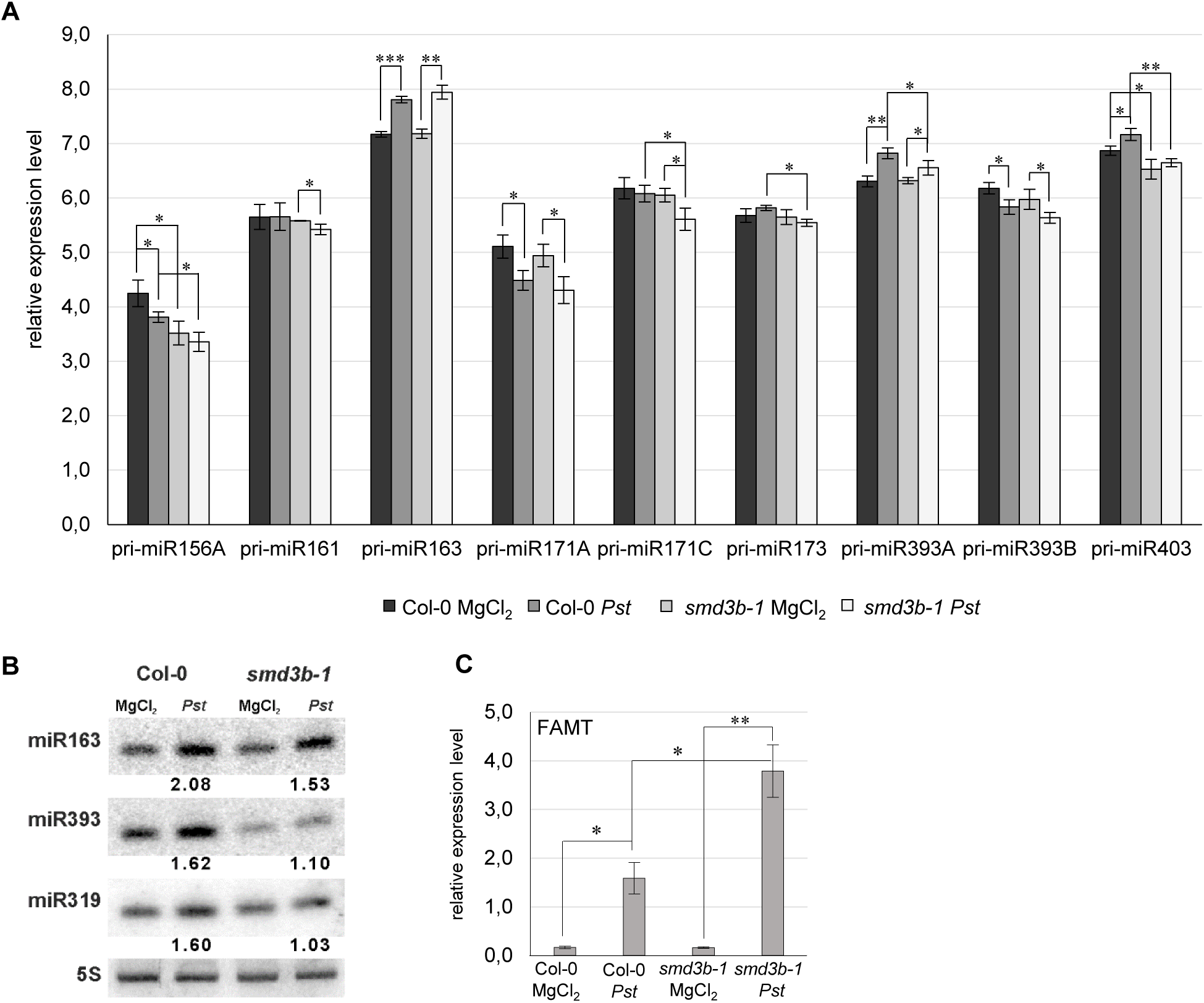
Both *smd3b-1* mutation and *Pst* infection modify the level of pri-miRNA and mature miRNA. (a) RT-qPCR analysis of chosen pri-miRNAs during *Pst* infection. Samples were collected from 6-week-old control (MgCl_2_) and infected (48 hpi, *Pst*) Col-0 and *smd3b-1* plants. Results represent mean of three independent biological replicates with error bars showing *SD*; \**P* < 0.05; \*\**P* < 0.01; \*\*\**P* < 0.001 (Student’s t-test). *GAPDH* mRNA was used as a reference. (b) Northern blot analysis of miRNAs involved in pathogen response. Numbers represent the levels of miRNA in *Pst*-treated Col-0 and the *smd3b-1* mutant relative to control (MgCl_2_), normalized to 5S rRNA loading control. Experiments were repeated at least three times; representative blots are shown. (c) RT-qPCR analysis of miR163 target, *FAMT* mRNA. Mean values ±SEM were obtained from three independent experiments; *P < 0.05; ** P < 0.01 (Student’s test). *UBC9* mRNA was used as a reference.

## DISCUSSION

Pre-mRNA splicing, especially alternative splicing, regulates many different cellular and physiological processes in plants, such as development, signal transduction and response to environmental cues, including biotic stress caused by microbial attack (Meyer *et al*., 2015; Yang *et al*., 2014). Still, despite realization that the majority of expressed genes in *Arabidopsis* undergo alternative splicing upon *P. syringae* infection (Howard *et al*., 2013), the extent of this level of regulation has not been extensively evaluated. We assessed long-term effects of splicing deficiency on plant immunity and we showed that a general splicing defect in a core spliceosomal component mutant, *smd3b-1*, results in decreased resistance to virulent *Pst* DC3000. Similar effects on pathogen proliferation were also observed for the *smd1b* mutant in another spliceosomal core protein, suggesting the involvement of the whole Sm complex. Our analyses reveal that spliceosome dysfunction impacts several aspects of pathogen response, namely stomatal immunity, activation of resistance-related factors and pathogen-associated WRKY transcription factors, ROS production and miRNA-dependent fine-tuning of plant defense (Figure 7). Our global transcriptome analysis showed that *smd3b-1* mutation affects mRNA levels and splicing pattern both in normal conditions and during *Pst* infection. Our data also adds to the description of transcription- and splicing-mediated reprogramming of gene expression caused by pathogen-induced stress in plants.

**Figure 7.**
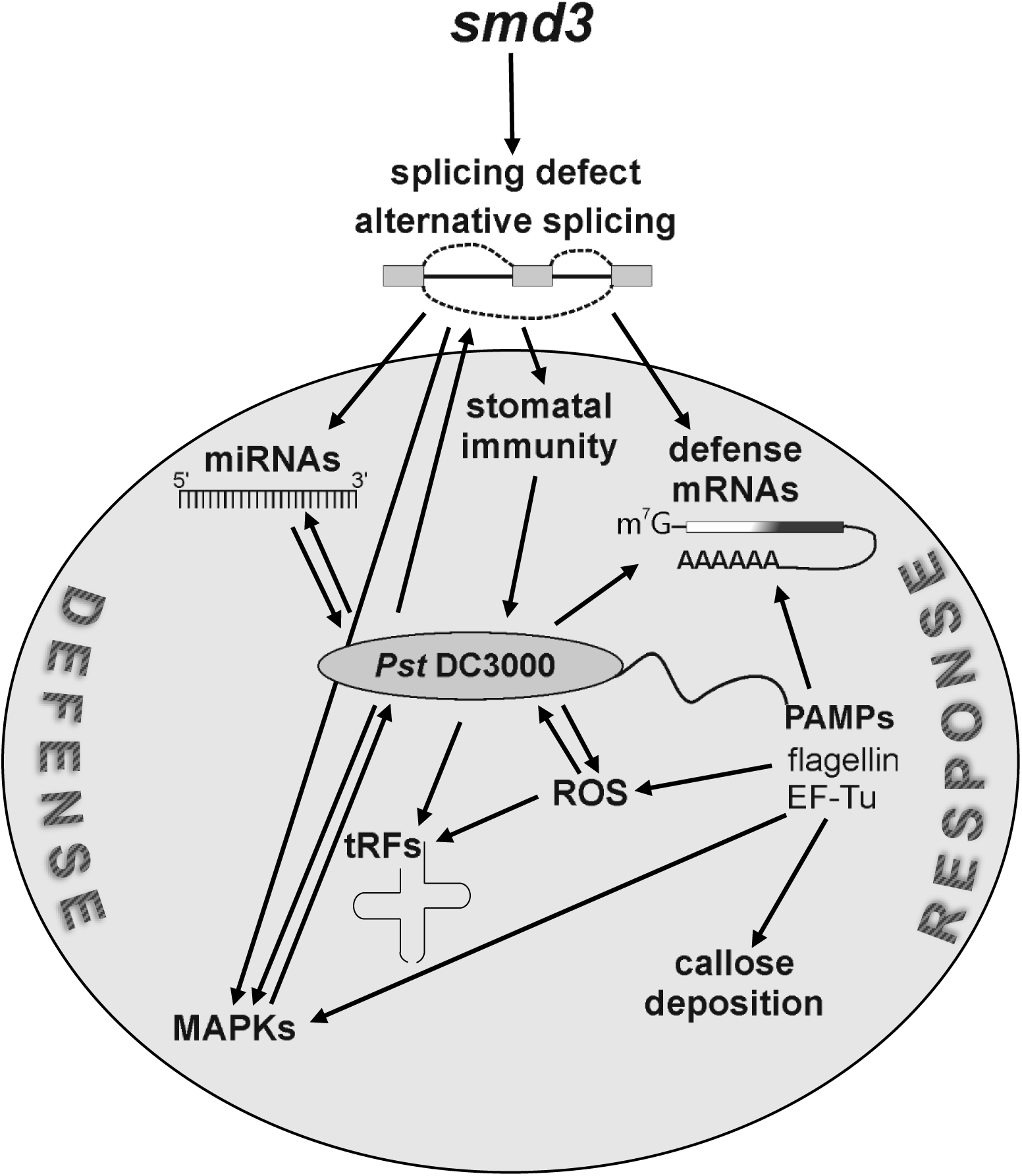
SmD3 affects several aspects of plant immunity through regulation of splicing of key pathogenesis factor mRNAs. Splicing defects in *smd3* mutants impact induction of pathogen-related proteins, including WRKY transcription factors, MAPKs cascade signaling, ROS production and miRNA-dependent fine-tuning of plant defense.

### Transcriptome upon *Pst* infection shapes *smd3b-1* defense response

Recent high-throughput RNA-seq analyses of transcriptome dynamics in *Arabidopsis* plants following infection with virulent DC3000 or ETI-triggering avirulent *Pst* strains (AvrRpt2 and AvrRpm1) showed that transcriptional response to avirulent pathogens was really fast, already observed at 4 hpi, whereas the equivalent response to virulent *Pst* was much slower and reached the same level at 24 hpi (Mine *et al*., 2018). We focused on the long-term response (48 hpi) to virulent *Pst* in Col-0 and the *smd3b-1* mutant to assess changes resulting from splicing defects. In line with previous reports we observed upregulated expression of many genes that belong to GO terms related to defense response and downregulation of chloroplast-associated genes (Lewis *et al*., 2015; Mine *et al*., 2018; de Torres Zabala *et al*., 2015). Changes in these ontology categories are characteristic of pathogen response as they include many genes that are key regulatory factors of plant immunity. We also note that, as reported, the number of splicing events and genes with splicing events were increased following pathogen treatment (Howard *et al*., 2013). The enhancement of alternative splicing associated with pathogenic infection further underlines the correlation between these two processes. Transcriptome profiles of *smd3b-1* control and *Pst*-treated plants revealed a complex picture of fluctuations in the expression pattern of genes related to different aspects of plant immunity (see below). Among these the most spectacular is a strong downregulation of a number of genes encoding key *PR* factors in the mutant in control conditions as well as enhanced activation of pathogenesis markers following bacterial infection. Plants lacking SmD3-b also exhibit altered expression of pathogenesis regulatory transcription factors as well as components of the BIK1 degradation pathway. In addition, our RNA-seq data revealed that *smd3b-1* mutation affected the splicing pattern of several of pathogen response-related factors. We envisage that the resultant of these changes, including variation in AS events, may contribute to dysregulated response to pathogen and its effectors in the *smd3b-1* mutant.

Although regulation of plant immunity in *Arabidopsis* by pre-mRNA splicing has been reported for several splicing factors (Meyer *et al*., 2015; Yang *et al*., 2014), our analyses present evidence supporting such a role for the core spliceosome. Sm proteins interact physically and functionally with pICln and PRMT5 components of the methylosome complex that mediates snRNP assembly, and the spliceosome activating nineteen complex (NTC) (Deng *et al*., 2016). Considering that these factors act as negative and positive regulators of plant immunity, respectively (Huang *et al*., 2016; Monaghan *et al*., 2009; Monaghan *et al*., 2010; Palma *et al*., 2007; Xu *et al*., 2012; Xu *et al*., 2011), their combined action in controlling disease resistance signaling via modulation of splicing is a strong possibility.

### Lack of SmD3-b impacts the pre-invasive stage of defense response and may lead to enhanced SAR and defense priming

Stomata are an integral part of the plant immune system and regulation of their aperture prevents pathogen entry into leaves and subsequent colonization of host tissues and disease symptoms. The *smd3b-1* mutation results in altered expression of a whole set of genes involved in stomata development and movement (Dataset S3), including positive and negative regulators of stomatal density and patterning. Moreover, increased stomatal density and aperture in *smd3b-1* plants together with their sensitivity to pathogen delivered via stomata suggest that SmD3b dysfunction impacts mainly the pre-invasive stage of defense response.

Another aspect of bacterial propagation is related to stomatal dynamics during infection. Briefly, PAMP-triggered stomatal closure to restrict pathogen entry, followed by SA-dependent basal defense, are suppressed by *Pst* effectors and coronatine that activates the antagonistic JA pathway and leads to stomatal reopening (Geng *et al*., 2014; Luna *et al*., 2011; Melotto *et al*., 2008). Although expression of several genes responsible for stomatal movement is altered in *smd3b-1* plants, stomatal closure appears not be compromised, probably due to the opposing impact of the *smd3b-1* mutation on expression of these genes. On the other hand, *Pst*-induced stomatal reopening could be affected in plants lacking Smd3-b due to the enhanced activation of NAC transcription factors that are induced by COR, leading to the COR-mediated stomatal reopening and thus more effective pathogen penetration.

Interestingly, *smd3b-1* mutation may affect SAR and defense priming that protect uninfected parts of the plant against secondary infections by a broad spectrum of pathogens and activate a faster and more robust response (Ádám *et al*., 2018; David *et al*., 2019; Fu and Dong, 2013). First of all, genes involved in the synthesis and modification of SA and NHP (e.g. *EDS5, PBS3*, *SID2/ICS1, FMO1*, *ALD1, BSMT1* and *MES9/SABP2*) that are important regulators of SAR and defense priming are strongly induced by *Pst* infection in *smd3b-1*. Notably, the expression of some of these genes (e.g. *SID2/ICS1* and *PBS3*) is regulated by WRKY46, WRKY53 and WRKY70 transcription factors (Wang *et al*., 2006), which are also upregulated in the mutant following infection. In turn, the level of *NRT2* (*NITRATE TRANSPORTER 2*) after *Pst* treatment is markedly decreased in the mutant, but not in the wild-type, and this may lead to constitutive priming (Camañes *et al*., 2012). Finally, defense priming and SAR also depend on ROS generation and callose deposition (Conrath *et al*., 2015; Mauch-Mani *et al*., 2017), and these are enhanced in PAMP-treated mutant plants. These observations suggest that in the absence of SmD3-b both SAR and priming defense may be enhanced, possibly to counteract the compromised stomatal immunity.

### Photosynthesis and chloroplast-associated genes in *smd3b-1* plants

PAMP perception leads to a general suppression of nuclear encoded chloroplastic genes and inhibition of photosynthetic processes, leading to ROS burst and defense response (Lewis *et al*., 2015; Lu and Yao, 2018; Serrano *et al*., 2016; de Torres Zabala *et al*., 2015). Our clustering analysis revealed that these genes are indeed strongly downregulated in Col-0 in response to *Pst*. Their expression was often decreased by *smd3b-1* mutation alone and was not further modified by pathogen attack. This may alter downstream events in the response pathway. Consistently, production of photosynthesis-derived, chloroplastic ROS was more robust in plants lacking SmD3-b, probably resulting in a stronger induction of many defense response genes. On the other hand, the primary apoplastic ROS burst was less pronounced in the mutant, possibly as a result of RBOHD attenuation. Additional changes in the apoplastic oxidative burst could stem from deregulation of splicing of other factors involved in ROS production and signaling (Kadota *et al*., 2015; Qi *et al*., 2017; Waszczak *et al*., 2018). Indeed, in *smd3b-1* plants several genes, such as *BAK1*, *XLG2*, *PRX34*, *AGG1*, *CRK4* and *CRK6*, showed statistically significant changes in the pattern of AS events that in particular apply to a higher number of retained introns.

From our analyses of the impact of SmdD3-b dysfunction on plant defense emerges a pattern whereby the initial response, including compromised stomatal immunity and limited production of the PAMP-triggered apoplastic ROS, leads to increased susceptibility to bacterial infection. This is followed by changes aiming at reinforcing plant defense systems through a more robust production of chloroplastic ROS, intensified hormonal signaling, enhanced callose deposition and stronger activation of defense-related genes. The interplay between these elements results in a complex and often opposing output of the mutant defense response. Importantly, this behaviour accompanies surface inoculation of the pathogen that closely resembles a natural infection, and does not take place when bacteria are artificially infiltrated into the leaf intercellular space.

The *smd3b* mutant displays a range of physiological phenotypes, including impaired root growth, altered number of floral organs and late flowering (Swaraz *et al*., 2011). These phenotypes correlate well with extensive changes in gene expression and differences in the splicing pattern of a few hundred of pre-mRNAs. It is tempting to speculate whether there are connections between these morphological and molecular phenotypes and dysregulation of the response to bacterial pathogen, considering that similar effects were observed for several other *Arabidopsis* mutants with defects in pre-mRNA splicing. It is possible that defective pre-mRNA splicing impact general fitness of the plant, which becomes more prone to disease progression, but based on growing evidence it seems more likely that splicing and alternative splicing are involved in regulation of pathogen response via adjusting the expression of key pathogenesis-related genes. We postulate that SMD3-b plays an important role not only in pre-mRNA splicing and spliceosome assembly but also acts as an intricate regulator of the plant defense response.

## EXPERIMENTAL PROCEDURES

### Plant material and growth conditions

*Arabidopsis thaliana* wild-type ecotype Columbia (Col-0) and *smd3a, smd3b* and *smd1b* mutant plants were used in this study: *smd3b-1* (SALK_006410) was a kind gift of Yoonkang Hur (Chungnam National University, Republic of Korea) (Swaraz *et al*., 2011); *smd1b* was received from Herve Vaucheret (INRA, CNRS, France) (Elvira-Matelot *et al*., 2016); *smd3b-2* (SALK_000746), *smd3a-1* (SALK_025193) and *smd3a-2* (SALK_020988) were purchased from NASC. Seeds were surface sterilized and grown for 2 weeks on MS medium (Murashige and Skoog, 1962) supplemented with 1% (w/v) sucrose and 0.3% phytogel under 16 h light/8 h dark photoperiod at 22/19°C. Infection experiments were performed on 6-week-old plants grown in soil under an 8 h light/16 h dark photoperiod at 22/19°C.

### Bacterial infection assays and PAMP treatments

Bacterial infection assays were performed with *Pseudomonas syringae* pv. *tomato* strain DC3000 (*Pst*) with density adjusted to 10^6^ cfu ml^-1^. 6-week-old plants were inoculated by spraying with *Pst* suspension in MgCl_2_/0.05% Silwet L-77 or with 10 mM MgCl_2_/0.05% Silwet L-77 (control). Material was harvested from at least 10 plants for each time point and used for RNA extraction. Bacterial growth was quantified as the number of dividing bacterial cells 24 and 72 hpi. Samples (four leaf discs) were taken from 2 leaves per six plants in each independent replicate.

For PAMP assays five days after stratification seedlings were transferred to 24-well plate with liquid MS (two seedlings per well). MS was exchanged for a fresh medium after 8 days and the next day flg22 (Alpha Diagnostic International Inc.), elf18 (synthetized by GL Biochem Ltd, Shanghai, China) or coronatine (Sigma) solution was added to a final concentration of 100 nM. Seedlings were harvested at the indicated time points.

### RNA methods

Total RNA was isolated from 2-week-old seedlings or 6-week-old plants using Trizol (Sigma) according to manufacturer’s instructions. Low-molecular weight RNAs were separated on 8% or 15% acrylamide/7 M urea gels and transferred to a Hybond N^+^ membrane (GE Healthcare) by electrotransfer. High-molecular-weight RNAs were analysed on 1.1% agarose/6% formaldehyde gels and transferred to a Hybond N^+^ membrane by capillary elution. Northern blots were carried out using γ-^32^P 5’-end-labelled oligonucleotide probes or random primed probes prepared using α-ATP^32^ and the DECAprimeTM II labelling kit (ThermoFisher Scientific). Quantification of northern blots was performed using Storm 860 PhosphorImager (GE Healthcare) and ImageQuant software (Molecular Dynamics). Oligonucleotides used for northern hybridization and RT-qPCR are listed in Table S2.

### Real-time PCR

RT-qPCR was carried out on cDNA prepared with mix of Random Primers and oligo-d(T) primer and SuperScript III Reverse Transcriptase (ThermoFisher) from 2 µg of RNA following DNase I digestion (Turbo DNase, ThermoFisher). Quantitative PCR was performed using SYBR Green I Master Mix (Roche) and the LightCycler 480 (Roche). Results were normalized to *UBC9* (*At4g27960*) or *GAPDH* (*At1g13440*) mRNAs.

### RNA-seq

Samples for RNA-seq were collected 48 hpi and total RNA was isolated using Trizol. RNA quality was verified with the Bioanalyzer 2100 (Agilent). Libraries were prepared from three independent biological replicates using Illumina TruSeq Stranded Total RNA with Ribo-Zero Plant rRNA Removal (Plant Leaf) protocol including barcoding and were paired-end sequenced by OpenExome s.c. For details see Supporting Information. RNA-seq data have been deposited in the Gene Expression Omnibus database under Accession Number GSE117077.

### Callose deposition assay

Sterilized Col-0 and *smd3b* seeds were sown in 6-well plates, containing MS medium and grown under long-day conditions for 7 days, when the medium was replaced by fresh MS. Plants were treated with flg22, elf18, and coronatine as described (Luna *et al*., 2011) at the final concentration of 1 µM for 24 h.

### ROS measurement

ROS production was detected using GloMax®-Multi^+^ Detection System (Promega) according to published protocols with minor modifications (Bisceglia *et al*., 2015; Smith and Heese, 2014).

## ACKNOWLEDGMENTS

We thank Yoonkang Hur (Chungnam National University, Republic of Korea) for *smd3b* seeds; Herve Vaucheret (INRA, CNRS, France) for *smd1b* seeds; Aleksander Chlebowski (IBB PAS) for assistance with microscopy and Wojciech Gierlikowski (Medical University of Warsaw) for assistance with ROS measurements.

## AUTHOR CONTRIBUTIONS

A.G. designed and performed most of the experiments; M.K. analysed RNA-seq data; M.K., M.S., J.P, J.D performed some of the experiments; A.J., Z.Sz.-K. supervised J.D.; A.G. conceived the project and wrote the manuscript with M.K. contribution; J.K. supervised and completed the writing.

## CONFLICT OF INTERST

The authors declare no conflict of interest.

## DATA AVAILABILITY STATEMENT

All relevant data are within the paper and its Supporting Information files.

## SUPPORTING INFORMATION

Additional Supporting Information may be found online in the Supporting Information section.

**Figure S1.**
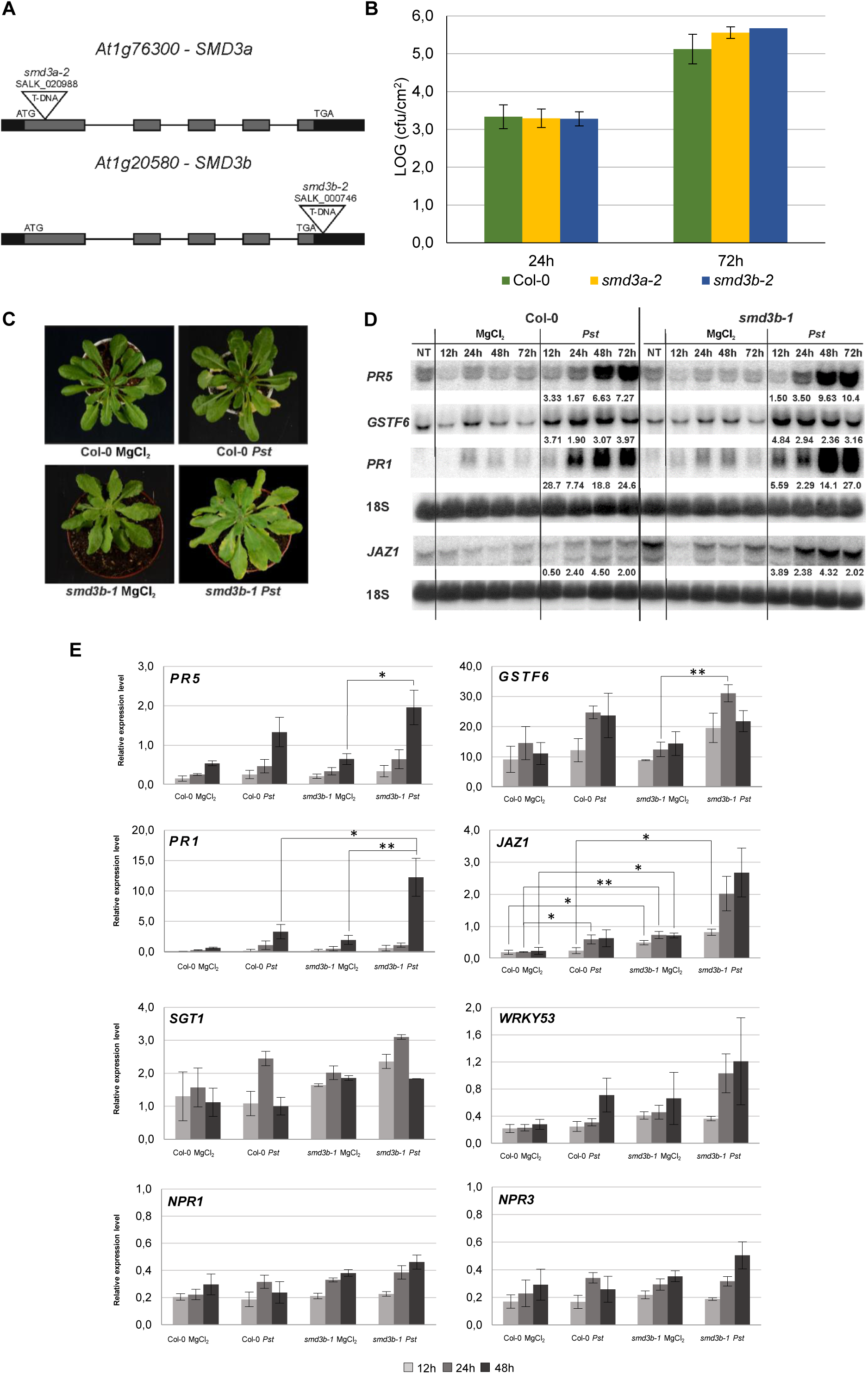
The *smd3b* and *smd3a* mutations cause changes in response to infection. (a) Structure of the *AtSMD3-a* (*At1g76300*) and *AtSMD3-B* (*At1g20580*) gene. Exons are represented by grey bars, UTRs are illustrated by black bars and localization of T-DNA insertions are indicated. (b) Growth of *Pst* DC3000 after 24 and 72 hpi in Col-0, *smd3a-2* and *smd3b-2* mutants. For each time point leaf discs were collected from 5 plants. Results are mean of two independent experiments. (c) Disease symptoms in Col-0 and *smd3b-1* 6-week-old plants (72 hpi). Experiments were repeated at least four times; representative pictures are shown. (d) Northern blot analysis of factors involved in pathogen response (another biological replicate). Samples were collected from non-treated (NT), control (MgCl_2_) and infected (*Pst*) Col-0 and *smd3b-1* plants at indicated time points. Numbers represent transcript level in *Pst*-treated Col-0 and the *smd3b-1* relative to control and normalized to 18S rRNA loading control. (e) RT-qPCR analysis of selected genes involved in pathogen response. Mean values ±SEM were obtained from three independent experiments; \**P* < 0.05; \*\**P* < 0.01 (Student’s t-test). *UBC9* mRNA was used as a reference.

**Figure S2.**
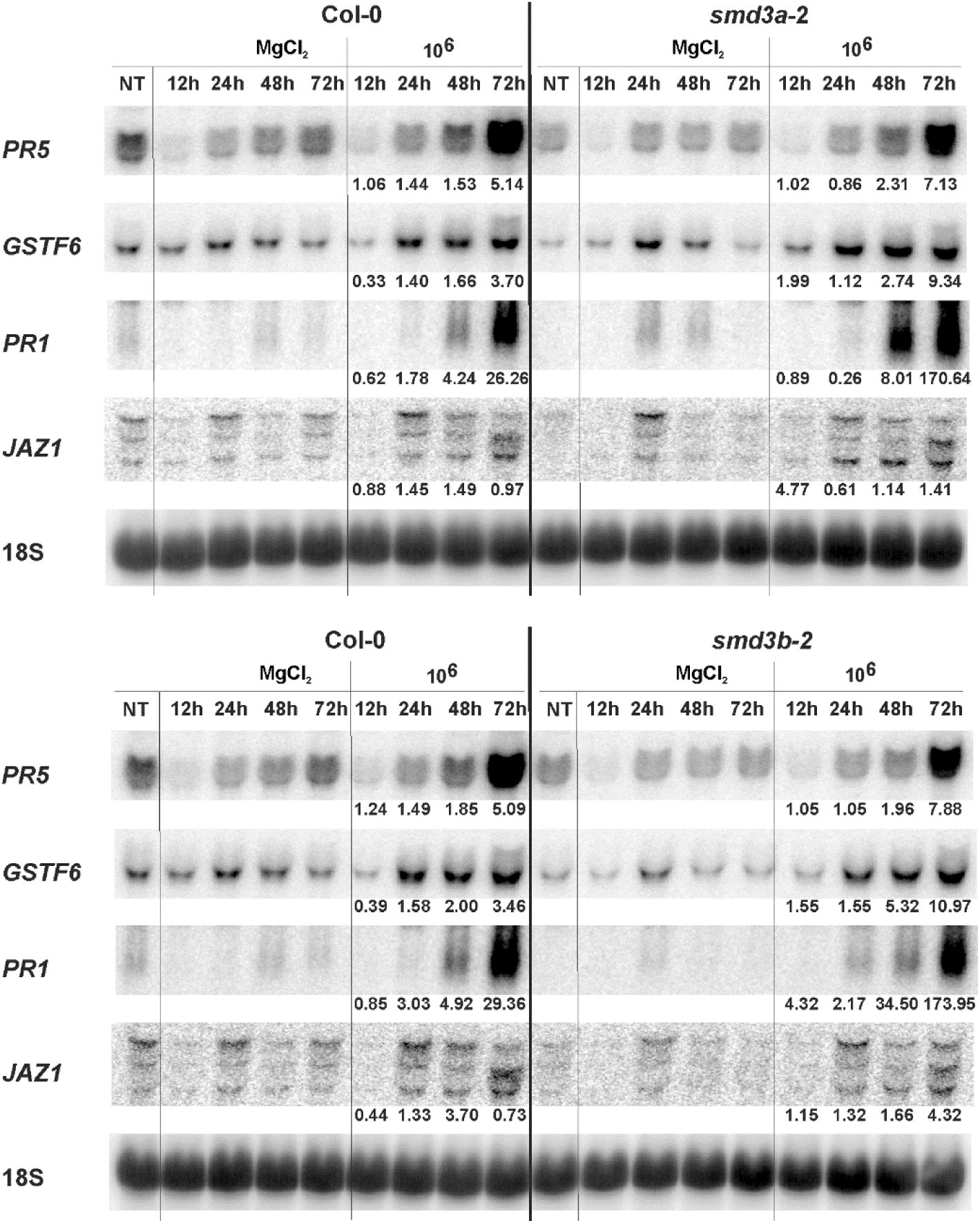
*Pst*-induced expression of pathogenesis markers in the *smd3a-2* and *smd3b-2* mutants. Northern blot analysis of factors involved in pathogen response. Samples were collected from non-treated (NT), control (MgCl_2_) and infected (*Pst*) Col-0, *smd3a-2* and *smd3b*-2 plants at indicated time points. Numbers represent transcript level in *Pst*-treated Col-0, *smd3a-2* and *smd3b*-2 relative to control and normalized to 18S rRNA loading control.

**Figure S3.**
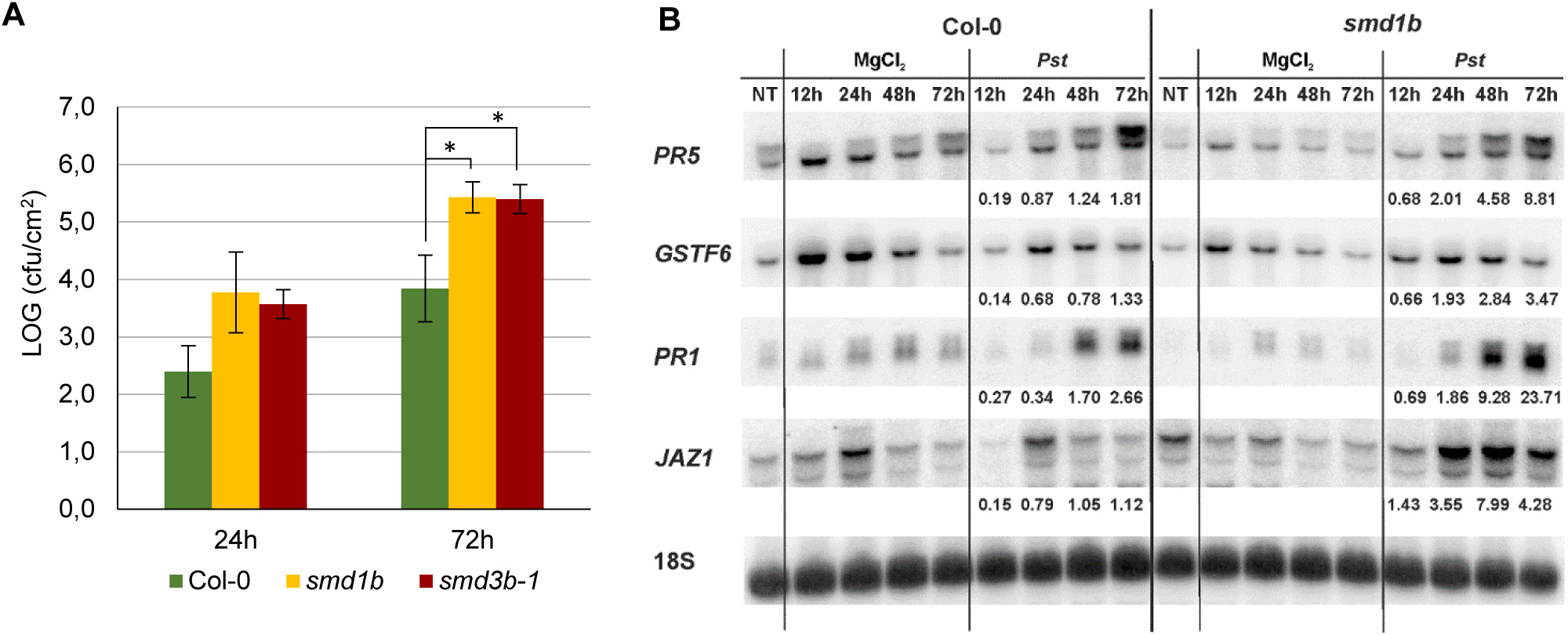
Both *smd1b* and *smd3b-1* mutants are susceptible to *Pst* infection. (a) Growth of *Pst* DC3000 after 24 and 72 hpi in Col-0, *smd1b* and *smd3b-1* mutants. For each time point leaf discs were collected from 5 plants. Results are mean of two independent experiments and error bars represent SD; *P < 0.05; ** P < 0.01 (Student’s t-test). (b) Northern blot analysis of factors involved in pathogen response. Samples were collected from non-treated (NT), control (MgCl_2_) and infected (*Pst*) Col-0 and *smd1b* plants at indicated time points. Numbers represent the ratio of transcript level in *Pst-*treated Col-0 and *smd1b* relative to control and normalized to 18S rRNA loading control.

**Figure S4.**
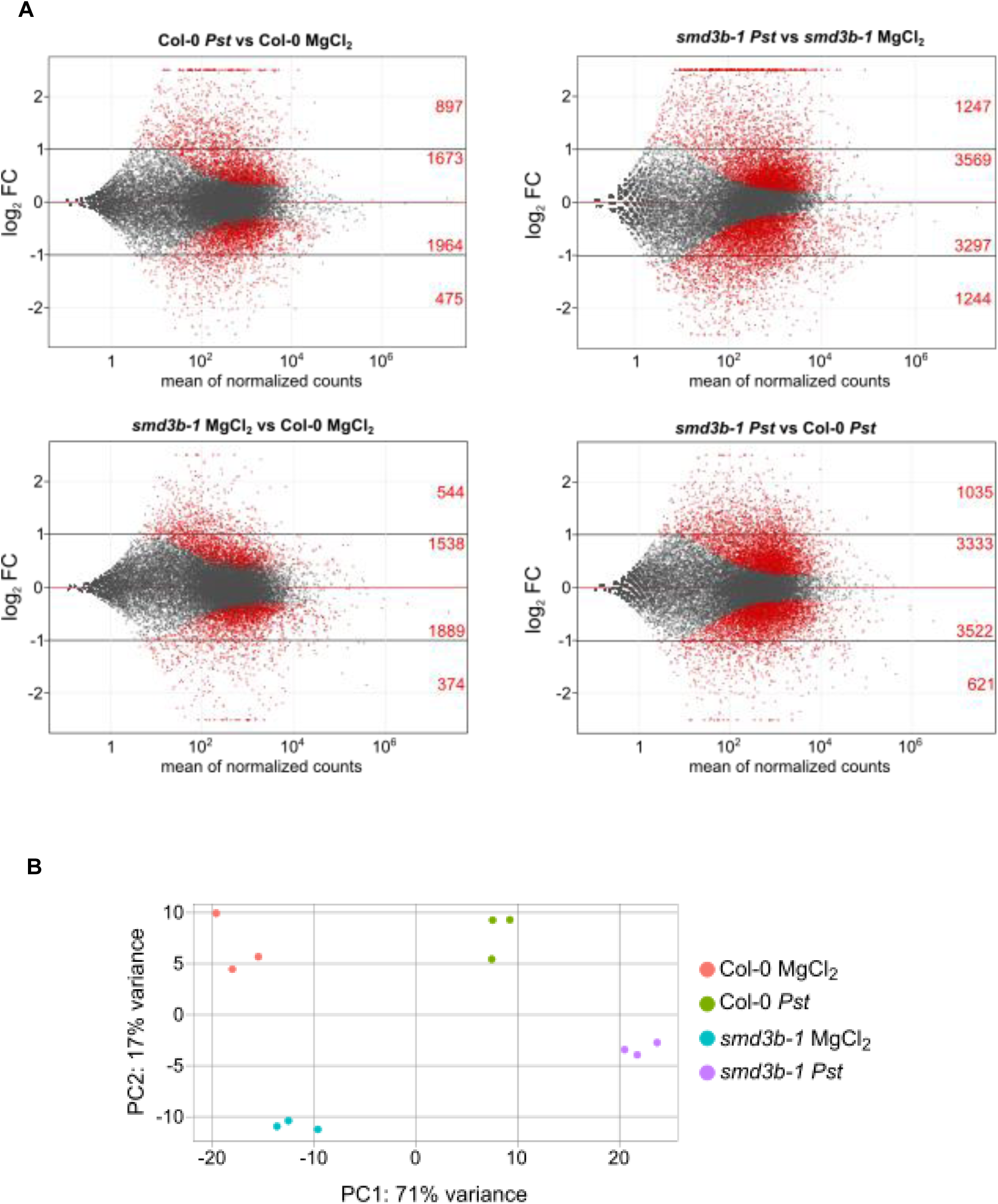
Results of RNA-seq analysis. (a) MA-plots (log_2_FC as a function of mean expression level on a logarithmic scale) of sequencing results for different comparisons, statistically significant hits (FDR < 0.05) are shown in red. Red numbers denote genes with significantly changed expression in each subgroup: up- or down-regulated, with absolute log_2_FC < 1 or >1. (b) PCA (Principal component analysis) shows that biological replicas in RNA-seq create four groups based on the presence of the *smd3b-1* mutation and *Pst* infection.

**Figure S5.**
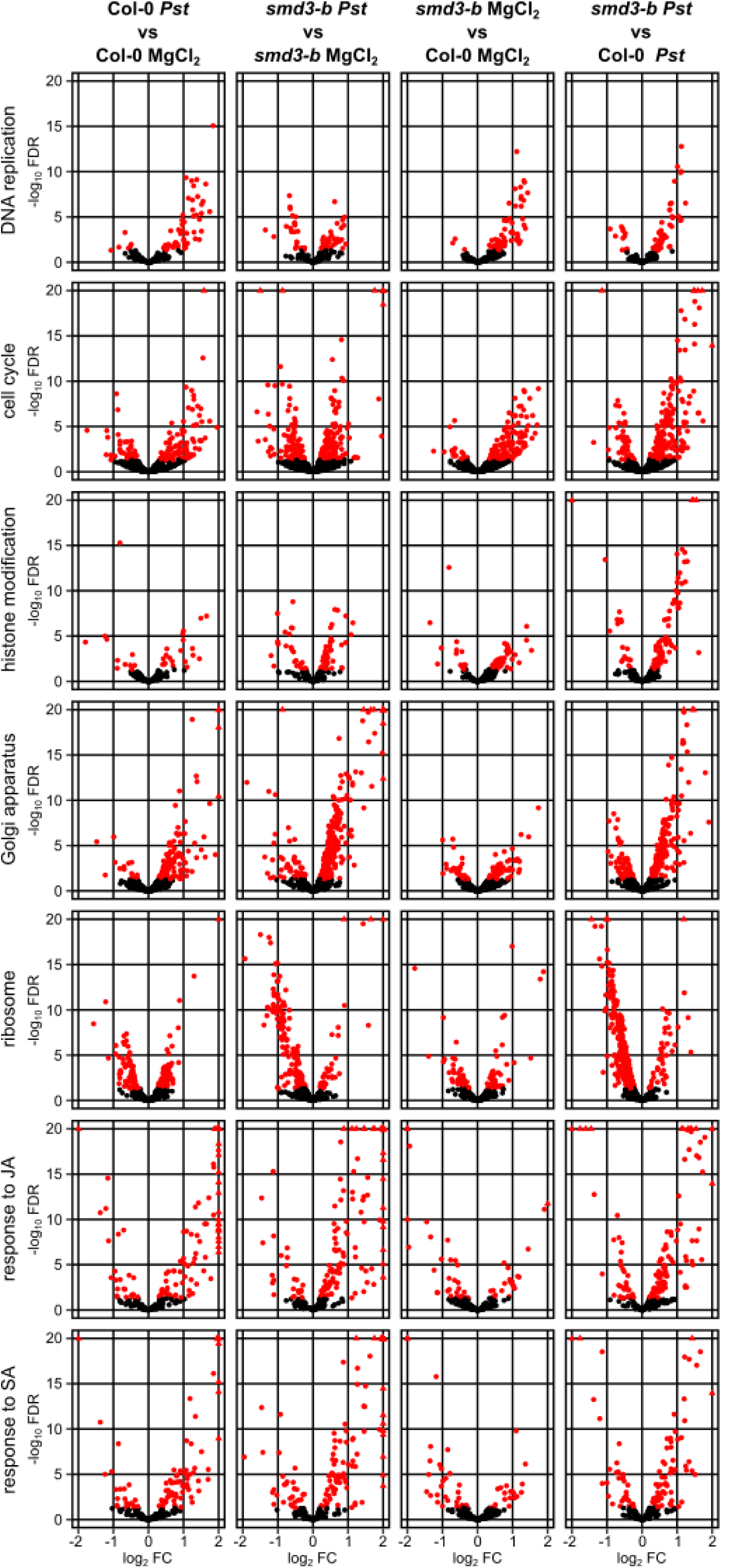
Both *Pst* infection and *smd3b-1* mutation impact mRNA expression. Bacterial infection and the *smd3b-1* mutation impact gene expression in selected categories. Volcano plots (-log_10_FDR as a function of log_2_FC) for genes implicated in DNA replication, cell cycle, histone modification, Golgi apparatus, response to jasmonic acid (JA) and salicylic acid (SA) and encoding components of ribosome. Red dots and triangles mark genes with significant changes.

**Figure S6.**
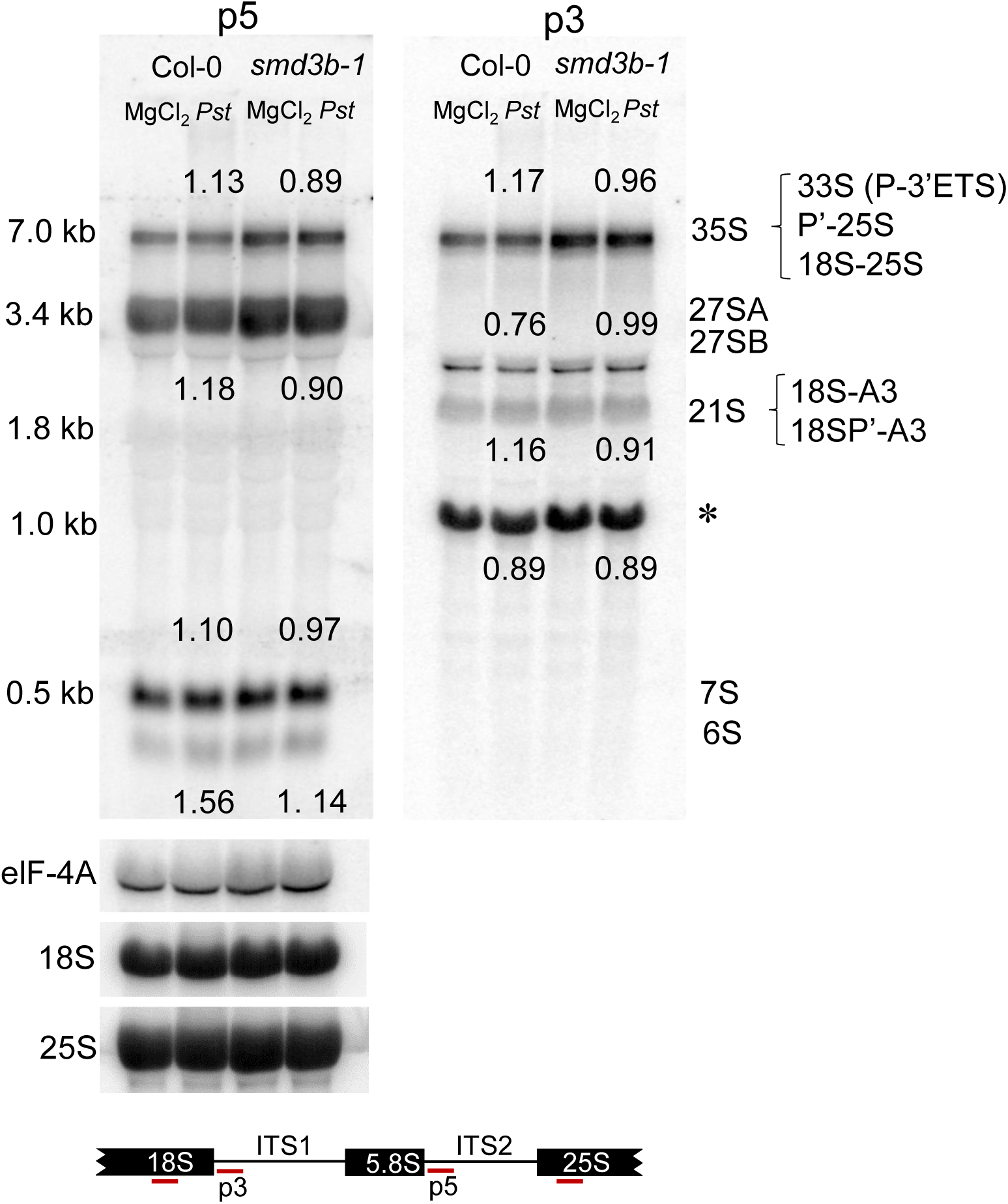
SmD3-b affects pre-rRNA processing. Northern blot analysis of rRNA precursors and intermediates in Col-0 and the *smd3b-1* mutant. Samples were collected from 6-week-old control (MgCl_2_) and infected (48 hpi, *Pst*) plants. rRNA precursors and intermediates are described on the right; molecular weight of 35S, 25S and 18S rRNA species are on the left. The position of specific *p5, p3,* 25S and 18S probes used for hybridization is shown in the diagram below. Numbers represent the ratio of the level of individual rRNA species in *Pst-*treated Col-0 and *smd3b-1* relative to control and normalized to *eIF-4A* mRNA loading control. Asterisk indicates cross-hybridization to organellar rRNA detected with probe *p3*.

**Figure S7.**
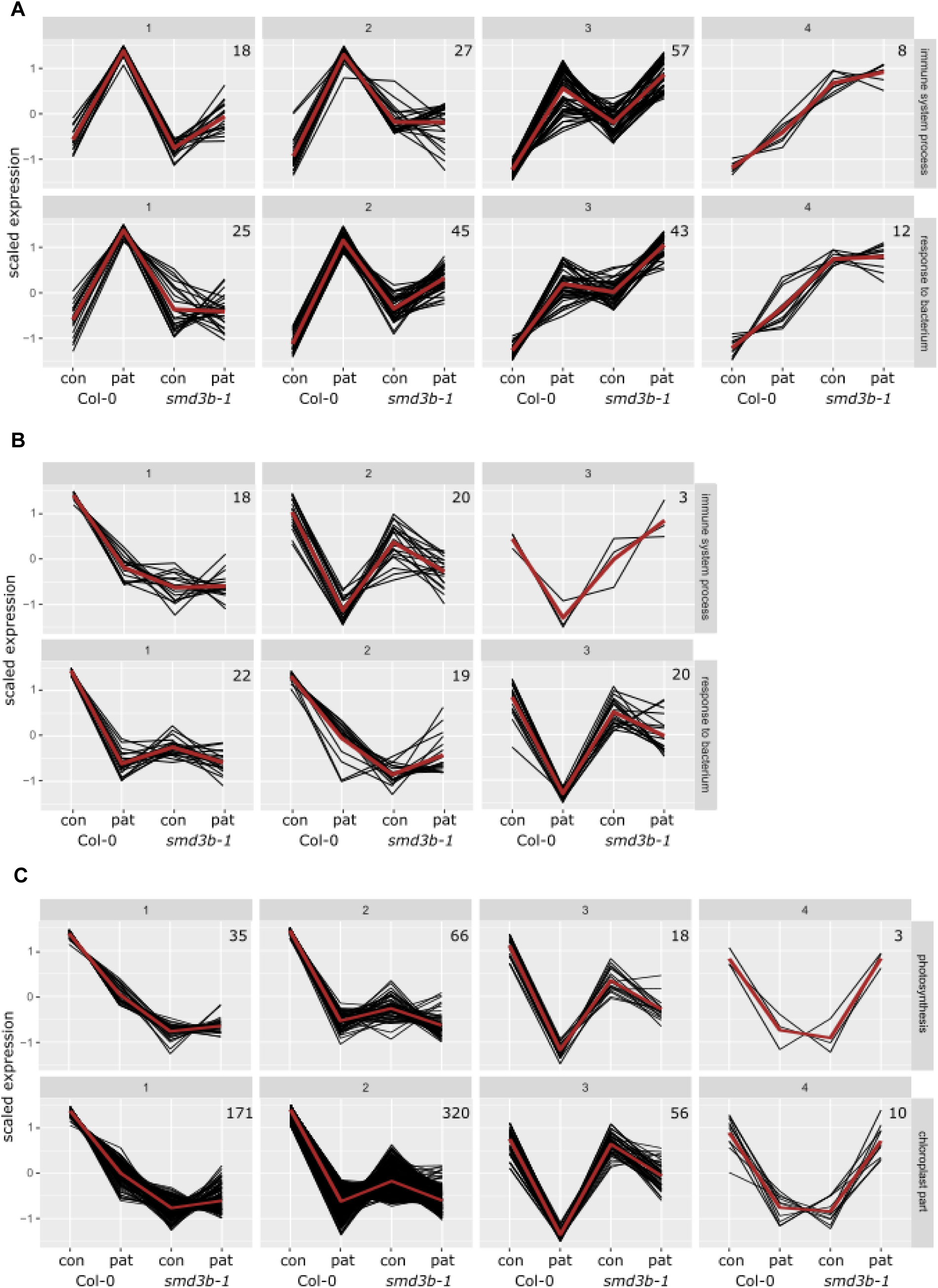
Differences between *smd3b-1* and Col-0 plants in expression of genes related to pathogenesis by clustering analysis. Clustering of gene expression profiles for genes affiliated with “response to bacterium”, “immune system process” (a, b), “photosynthesis” and “chloroplast part” (c) GO terms that were significantly upregulated and downregulated after pathogen treatment. Number of genes in each cluster are depicted on each diagram. Clustering was performed using standard R functions.

**Figure S8.**
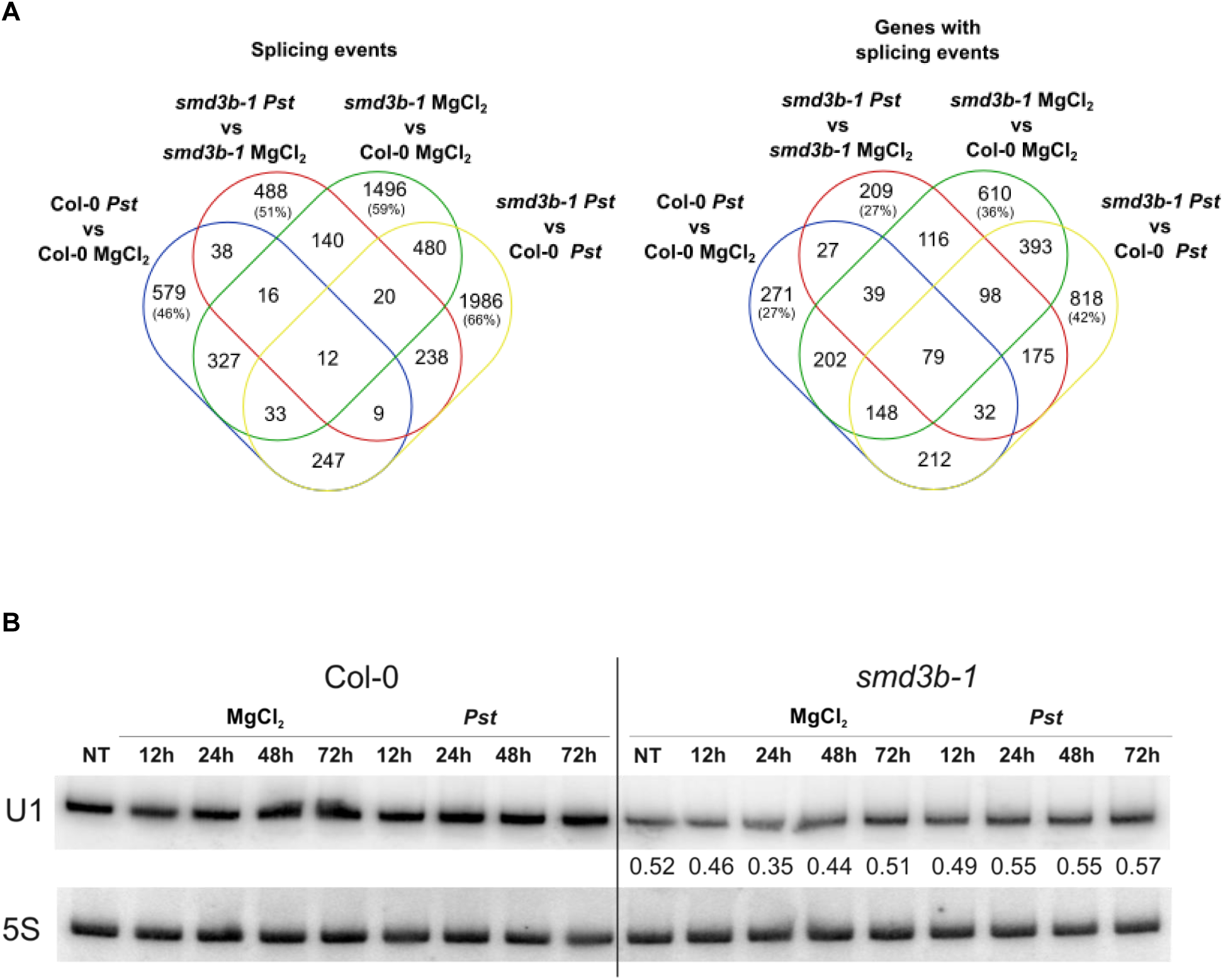
Splicing events and genes with splicing events significantly altered in Col-0 and the *smd3b-1* mutant upon treatments. (a) Venn diagrams indicating the number of significant overlapping splicing events and genes with splicing events in Col-0 and the *smd3b-1* mutant after treatments. Percent of unique splicing events and genes with splicing events when compared to other sets is given in parentheses. (b) *smd3b-1* mutation confers global splicing defect. U1 snRNA is destabilized in the absence of SmD3-b. Northern blot analysis of U1 snRNA. Samples were collected from non-treated (NT), control (MgCl_2_) and injected (*Pst*) Col-0 and *smd3b-1* plants at indicated time points. Numbers represent transcript level in Col-0 and *smd3b-1* relative to control and normalized to 5S rRNA loading control.

**Figure S9.**
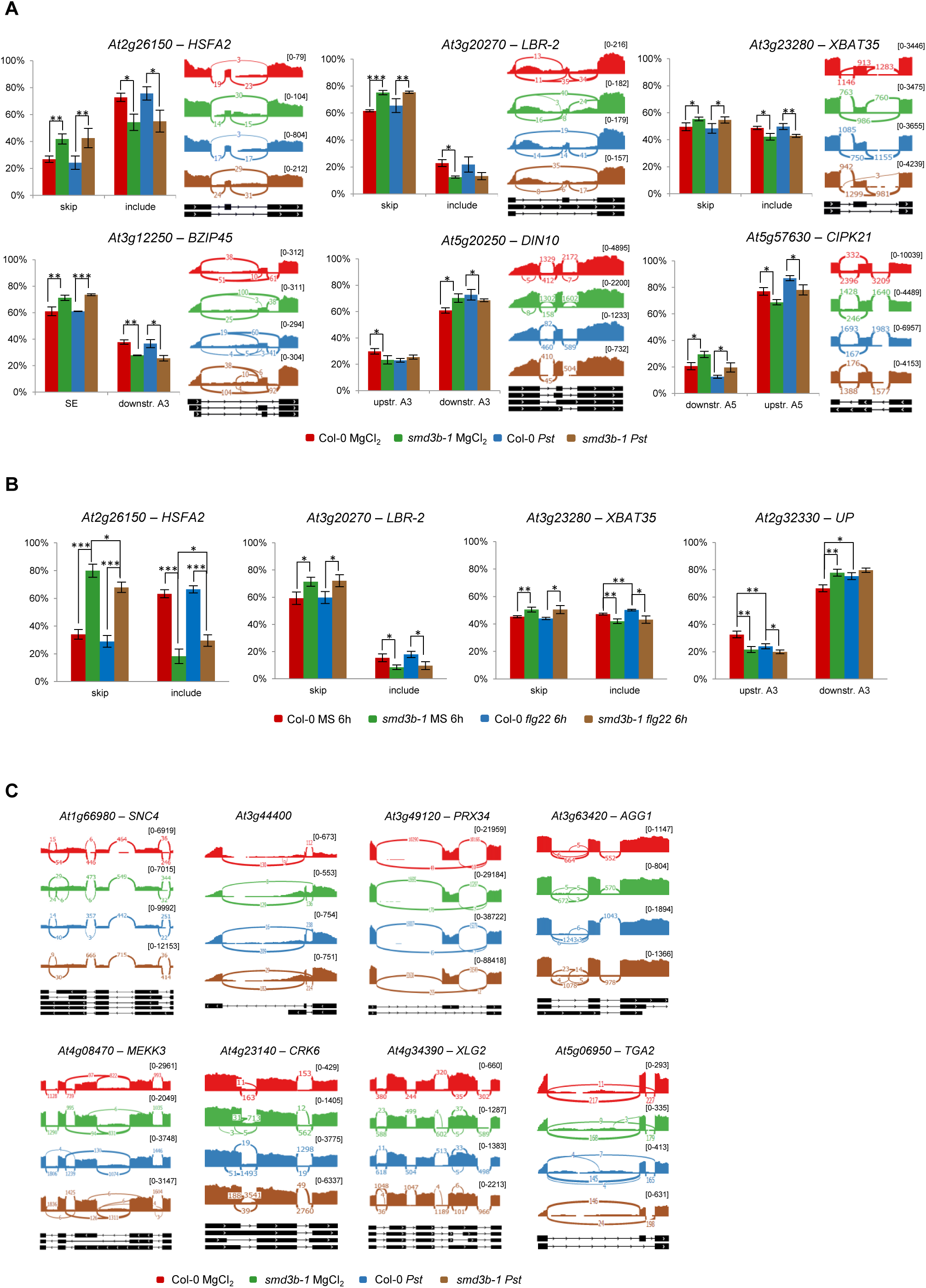
*Pst*, flg22 treatment and *smd3b-1* mutation affect alternative splicing (AS) events. (a) Analysis of AS events by RT-qPCR for selected genes in control (MgCl_2_) and infected (48 hpi; *Pst*) Col-0 and *smd3b-1* plants; the type of AS event is described below each graph: SE-skipped exon, RI-retained intron, A5/A3-alternative 5’ or 3’ splice sites, respectively. Results represents mean of three independent biological replicates with error bars showing *SD*; *P < 0.05; ** P < 0.01; \*\*\**P* < 0.001 (Student’s t-test). *GAPDH* mRNA was used as a reference. Sashimi plots were created from RNA-seq data using Integrative Genomics Viewer (IGV). The numbers in brackets are the range on the bar graph. Note differences in scales for each sashimi plot. (b) Analysis of AS events by RT-qPCR for selected genes in control-(MS) and flg22-treated (100nM for 6 h) Col-0 and *smd3b-1* plants. The type of AS event is described below each graph: SE-skipped exon, A3-alternative 3’ splice sites, respectively. Results represent mean of three independent biological replicates with error bars showing *SD*; *P < 0.05; ** P < 0.01; *** P < 0.001 (Student’s t-test). *GAPDH* mRNA was used as a reference. (c) Analysis of AS events for selected genes in control (MgCl_2_) and infected (48 hpi; *Pst*) Col-0 and *smd3b-1* plants. AS events are significantly altered for genes involved in pathogen response. Sashimi plots were created from RNA-seq data using Integrative Genomics Viewer (IGV). The numbers in brackets are the range on the bar graph. Note differences in scales for each sashimi plot.

**Figure S10.**
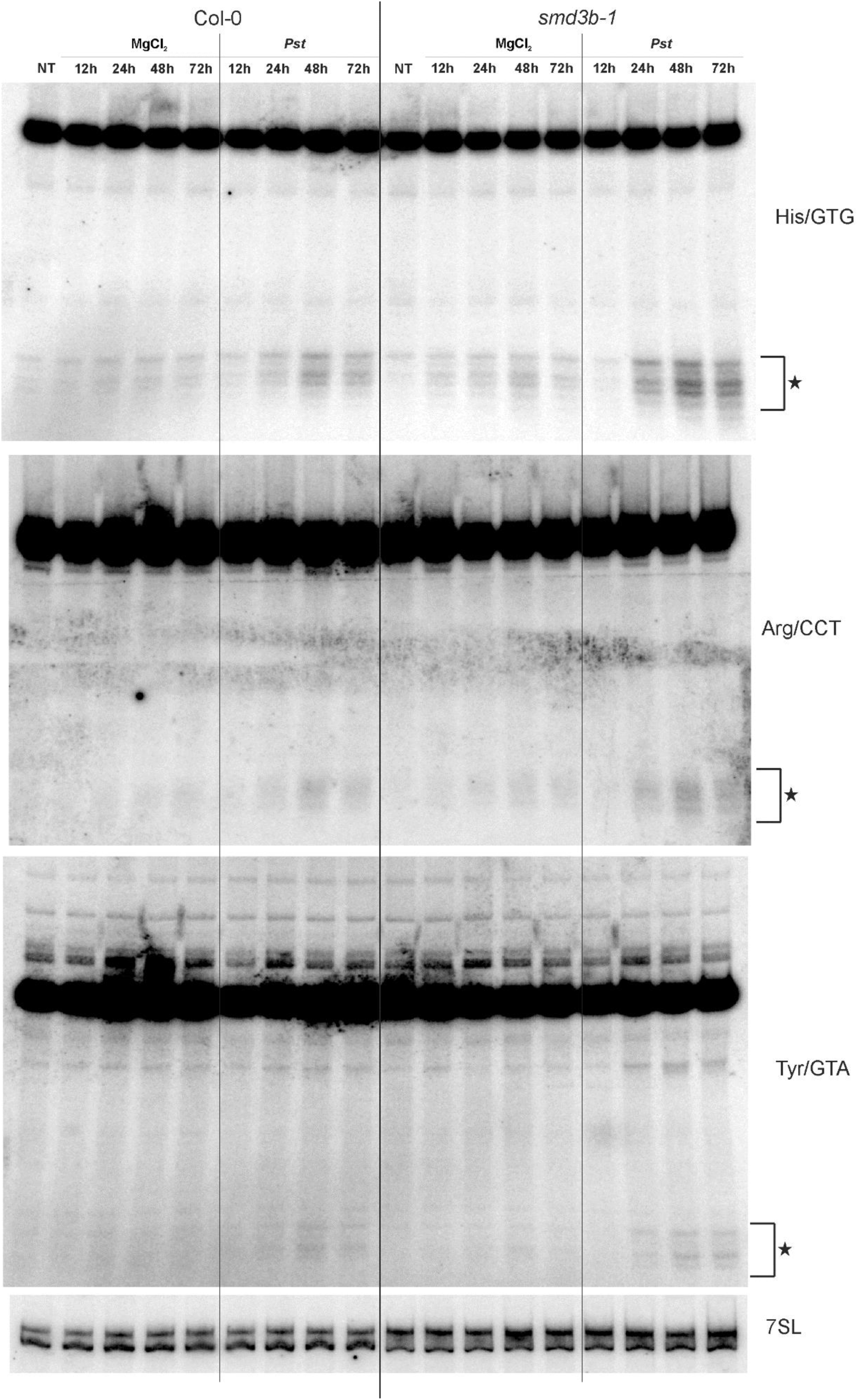
*Pst* DC3000 infection increases production of tRNA fragments in the *smd3b-1* mutant. Northern blot analysis (PAGE) of three tRNA species following the *Pst* treatment. Samples were collected from non-treated (NT), control (MgCl_2_) and infected (*Pst*) Col-0 and *smd3b-1* plants at indicated time points. 7SL was used as a loading control. Asterisks indicate accumulation of short RNA fragments (tRFs). Experiments were repeated twice with similar results; representative blots are shown.

**Figure S11.**
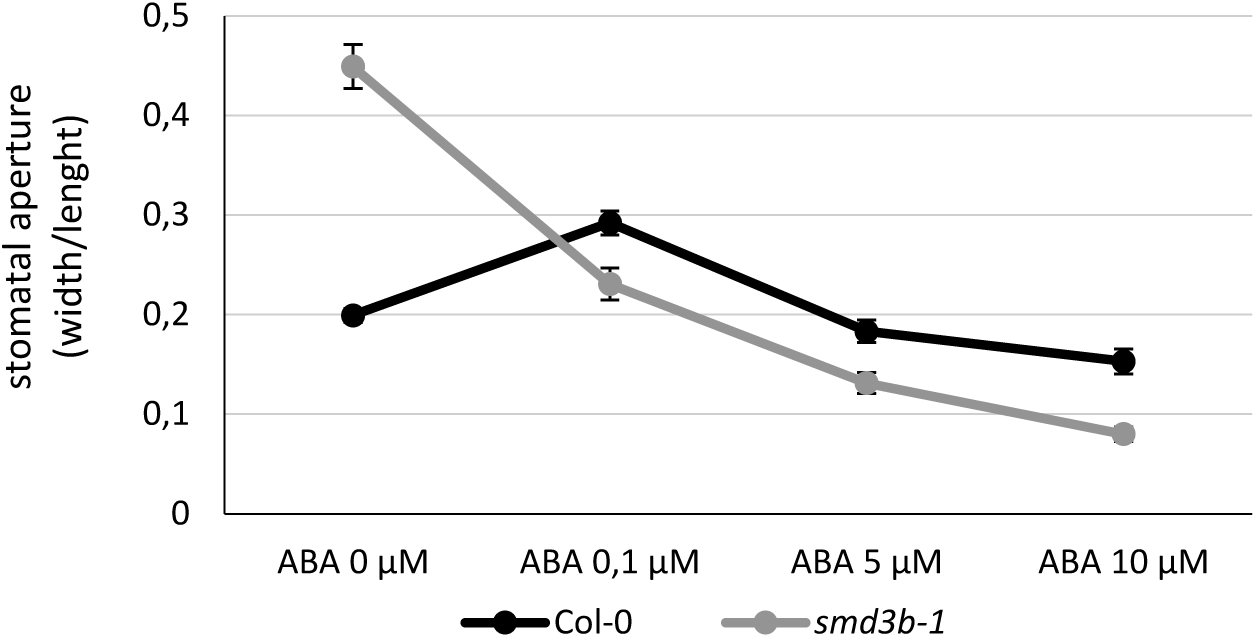
Stomatal closure following ABA treatment in Col-0 and *smd3b-1*. Mean values (±SEM) were obtained from six independent experiments with 40 stomata per data point; \**P* < 0.05; ***P<0.001 (Student’s t-test).

**Figure S12.**
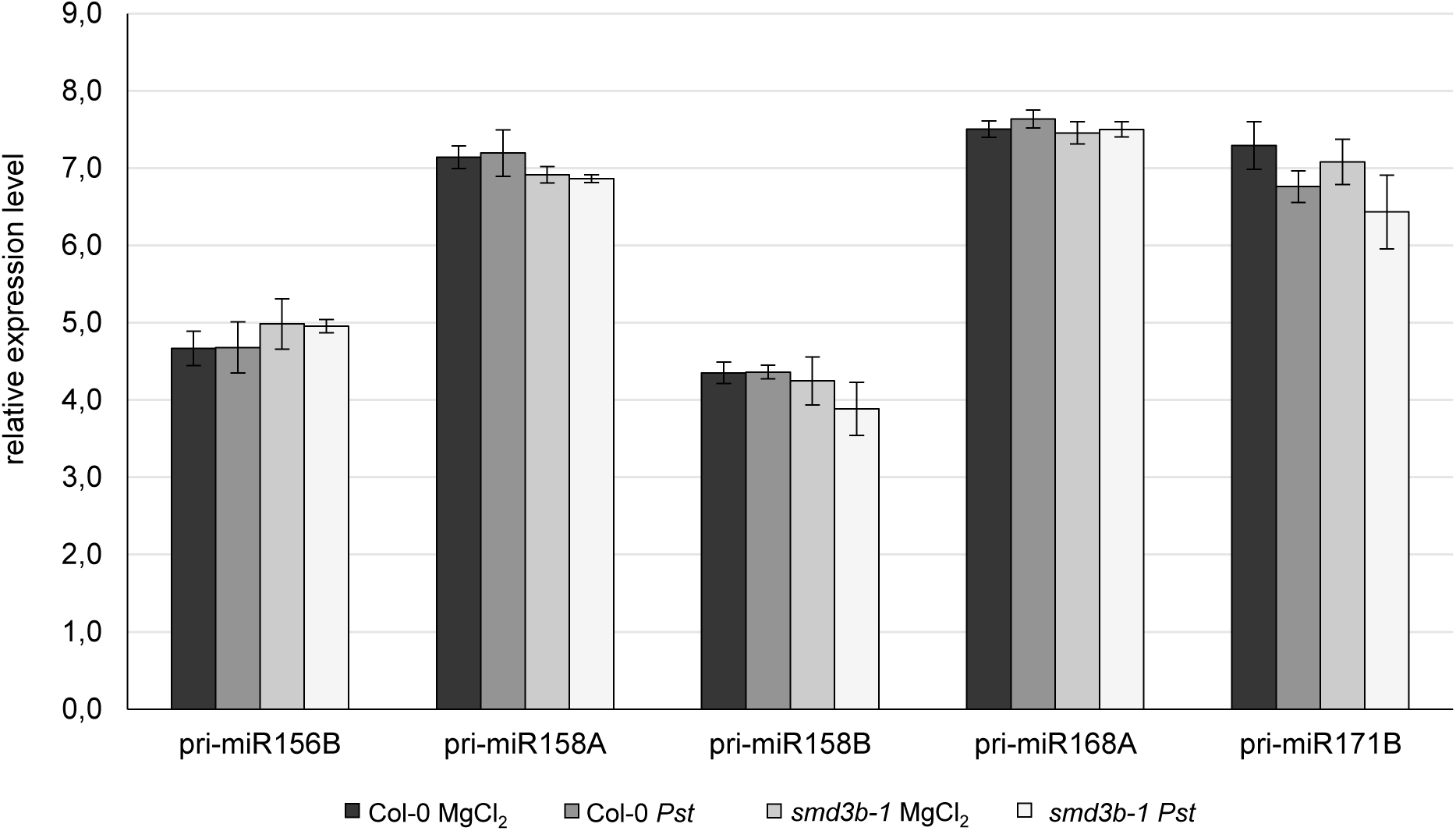
*Pst* infection changes the level of pri-miRNAs in the *smd3b-1* mutant. RT-qPCR analysis of chosen pri-miRNAs during *Pst* infection. Samples were collected from 6-week-old control (MgCl_2_) and infected (48 hpi, *Pst*) Col-0 and *smd3b-1* plants. Results represent mean of three independent biological replicates with error bars showing *SD* with no statistically significant changes. *GAPDH* mRNA was used as a reference.

**Table S1. Characteristic of genes involved in pathogen response having significantly affected alternative splicing events in the *smd3b-1* plants.**

**Table S2. List of primers used in this study.**

**Dataset S1. List of significantly affected genes based on RNA-seq.**

**Dataset S2. Significantly enriched Gene Ontology terms among different gene sets.**

**Dataset S3. List of genes involved in different aspect of pathogen response with significantly changed expression in the *smd3b-1* mutant.**

**Dataset S4. List of alternative splicing events.**

## SUPPORTING INFORMATION – Materials and Methods

### RNA-seq

The quality of the data was assessed using *fastqc* (v0.11.2 http://www.bioinformatics.babraham.ac.uk/projects/fastqc/). Reads for each sample were aligned to the TAIR10 *A. thaliana* genome from ensemble (release v29; (Kersey *et al*., 2016)) and reference annotation AtRTD2 (release 19.04.2016; (Zhang *et al*., 2017)) using STAR (v2.5.0a, (Dobin *et al*., 2013)) with the following command-line parameters:

*STAR --runMode alignReads --sjdbOverhang 149 --sjdbGTFfile --readFilesCommand --sjdbInsertSave All --outFilterType BySJout --outFilterMultimapNmax 10 --alignSJoverhangMin 10 --outFilterMismatchNmax 10 --alignIntronMax 100000 --alignMatesGapMax 100000 --outSAMattrIHstart 0 --outMultimapperOrder Random --outSJfilterIntronMaxVsReadN 5000 10000 15000 20000 --outSAMtype BAM SortedByCoordinate --quantMode GeneCounts --outSAMunmapped Within --sjdbFileChrStartEnd*

RNA-seq alignments were split to separate read-pairs that originate from transcription on the forward and reverse strands using samtools (v1.1; (Li *et al*., 2009)). Coverage graphs were calculated with *genomeCoverageBed* from *bedtools* (v2.17.0; (Quinlan and Hall, 2010)) with normalization to number of reads and were converted to bigwig format with bedGraphToBigWig (v4; http://hgdownload.cse.ucsc.edu/admin/exe/linux.x86_64/). Differential expression (DE) was performed using *DESeq2* (v1.16.1) *R* (v3.4.1) package with parameter *alpha = 0.05* (Love *et al*., 2014). Genes with FDR adjusted p-value <0.05 and absolute log_2_FC >1 were considered significantly changed. Clustering of gene expression profiles was performed using standard R functions on sets of genes selected based on their expression change and GO term affiliation. Splicing analysis was done using reference annotation AtRTD2 (release 19.04.2016; (Zhang *et al*., 2017)) and rMATS (v3.2.5; (Shen *et al*., 2014)) with command-line parameters: *-t paired-len 149-libType fr-firststrand-novelSS 1.* Differential splicing events with FDR < 0.05 and delta PSI > 0.05 were considered as significant. Sashimi plots were created using IGV from RNA-seq data (Thorvaldsdóttir *et al*., 2013).

### Analysis of stomatal density and aperture size

The experiments were performed using fully developed rosette leaves of 35-day-old *Arabidopsis* plants. The cleared epidermal peels from abaxial leaf surfaces and nail polish immersions from adaxial leaf surfaces were prepared and examined with a light microscope equipped with a Nikon Eclipse Ti camera, DS-Fi1c-U2 optics and Plan Apo VC 20x DIC N2. The counts were made on 6 leaves from independent plants for each ecotype and then averaged.

For the measurements of stomatal aperture preparations were made from epidermal peels from abaxial leaf surfaces. Before preparation the leaves were incubated in a buffer composed of 10 mM KCl, 0.1 mM EGTA and 10 mM MES-KOH at pH 6.15 for 2 hours. Measurements were analysed using a light microscope, Eclipse Ti camera, DS-Fi1c-U2 optics and Plan Apo VC 20x DIC N2 (Nikon,Tokyo, Japan). The ratio of the length and width of the pore between guard cells was calculated using the NIS-Element program. The counts were made on 6 leaves from independent plants.

### Bacterial infection assays by injection

Bacterial infection assays were performed with virulent *Pseudomonas syringae* pv. *tomato* strain DC3000 (*Pst*). Bacteria for inoculation were grown overnight in LB medium with rifampicin (50 μg ml^-1^) at 28°C, resuspended in 10 mM MgCl_2_ with density adjusted to 10^5^ cfu ml^-1^ (OD_600nm_ = 0.003). 6-week-old plants were inoculated by injection with *Pst* suspension in 10 mM MgCl_2_ and covered with plastic lids overnight. Material was harvested from at least 8 plants for each time point, frozen in liquid nitrogen and used for RNA extraction. Bacterial growth was quantified as the number of dividing bacterial cells 24 and 72 h after infection (hpi). Samples (four leaf discs) were taken using a cork-borer (4 mm) from 2 leaves per six plants in each independent replicate.

